# Facilitating Valid Statistical Inference in Biomedical Image Synthesis

**DOI:** 10.64898/2026.10.14.759239

**Authors:** Jiangmei Xiong, Shunxing Bao, Qi Liu, Ken S. Lau, Lori A. Coburn, Keith T. Wilson, Bennett A. Landman, Simon Vandekar, Siyuan Ma

## Abstract

Generative AI for medical image synthesis is increasingly used to impute missing data and augment biomedical datasets. Yet despite achieving high computational similarity to their real-world counterparts, synthetic images remain uncommon in clinical research practice, in part because of concerns about their biological validity. Here, we evaluate whether synthetic images support reliable downstream statistical analysis and biological inference in magnetic resonance imaging (MRI) and multiplexed immunofluorescence (mIF) imaging. We find that associations between patient phenotypes and synthesized images can be severely attenuated. This is because synthesis methods typically omit clinically relevant covariates and therefore generate images that, although visually realistic, mask key biological differences. Furthermore, naive analyses treat synthesized images as if they were observed, ignoring the statistical uncertainty introduced by synthesis. Drawing on multiple imputation theory, we introduce Phenotype Aware Multiple Synthesis (PAMS), an analysis framework that requires no model retraining and instead adjusts synthesized output to recover valid biological estimates with appropriately calibrated uncertainty. Applied to MRI and mIF data, PAMS reduced estimation bias and restored nominal confidence interval coverage. Because it is compatible with existing synthesis pipelines, PAMS offers a practical and rigorous route to statistically and biologically valid use of synthesized images in biomedical imaging research.

## 1 Introduction

Biomedical and pathology imaging, including magnetic resonance imaging (MRI) and multiplexed immunofluorescence (mIF), are indispensable components of biomedical research, providing qualitative and quantitative insights into biological samples [1–5]. These imaging modalities have been important in the advancement of research in cancer, neurological diseases, and inflammatory conditions, and their utility increases as quantitative image analysis methods become more sophisticated [6–10]. However, real world challenges in medical image acquisition, such as technical artifacts, modifications to protocols, or resource constraints are barriers to collecting adequate sample sizes of optimal quality, ultimately resulting in insufficient data sizes for desired scientific hypothesis or statistical power [11–17]. For example, head motion artifacts are prevalent in fMRI data and are both difficult to remove and costly to re-measure, leading to reduced sample sizes [18–20]. In response to similar challenges with data acquisition, biomedical image synthesis has emerged as an active area of research [21–29]. Image synthesis methods stemming from recent breakthroughs in deep learning and generative artificial intelligence (AI) have been used to generate uncollected channels or images, augment sparse datasets, increase image resolution, and recover lost imaging information [29–33].

Despite the improvement in the methodology and accuracy of image synthesis, the translational impact towards scientific and clinically relevant applications remains limited. So far, there are no well-established regulatory guidelines for the use of synthesized medical images, and the current use of synthesized images in the medical field is limited to education and training of machine learning models [33, 34]. Recognizing this gap, the U.S. Food and Drug Administration (FDA) Center for Devices and Radiological Health (CDRH) launched the Artificial Intelligence Program, and one of the primary research objectives is specifically aimed at validating and operationalizing the use of AI in supplementing medical patient datasets [35, 36]. In support of this objective, Zamzmi, G. et al. [37] highlighted concerns that synthesized data containing biologically unrealistic content may still achieve high computational similarity scores, and provided preliminary suggestions for establishing guidelines incorporating biological validity. Accordingly, these ongoing concerns about the validity, interpretability and generalizability of findings using synthesized data are reflected in the limited uptake of synthetic imaging in research [38–41]. Realizing the full potential of synthesized medical images in research and clinical workflows requires methods that explicitly ensure their biological and statistical validity, which is a prerequisite that current synthesis approaches do not fully address.

A critical yet underexplored dimension to ensure that biomedical image synthesis generates valid inference is the statistical calibration of bias (i.e., systematic deviation of the estimated quantity from truth) and uncertainty (i.e., variability of the estimated quantity given the images are synthesized rather than truth). Both are introduced by the synthesis process itself, including process error or failure to capture critical clinical information during training [42]. Recent machine learning research on the post-hoc correction of model-predicted values offers two solutions, each targeting one of these problems. Angelopoulos et al.[43] proposed prediction-powered inference, a framework that uses observed real data to “debias” estimations from synthesized data, with a primary focus on bias correction. Conformal prediction instead targets uncertainty, quantifying it in machine learning predictions [44, 45], and has been applied to spatial gene expression synthesis for spatial transcriptomics data with valid uncertainty estimation [46]. To this end, classical statistical missing data research offers an alternative solution that treats both problems jointly, namely, multiple imputation (MI) [47, 48]. Originally developed in the context of synthesizing missing census data, MI generates multiple realizations of synthesized data for analyses that (1) require key clinical variables be incorporated, and (2) properly reflect the uncertainty that the images are synthesized rather than truth. It thus addresses both bias and uncertainty [49, 50]. Under this context, multiple imputation can help provide a statistically rigorous basis for a broadly applicable method that accounts for such bias and uncertainty in synthesized images simultaneously.

To address these challenges in medical image synthesis, we introduce Phenotype Aware Multiple Synthesis (PAMS), a modular framework grounded in the multiple imputation paradigm that improves inferential validity regarding bias and uncertainty in the downstream analysis of synthesized images. To systematically assess the need for such a framework, we evaluated the biological and statistical validity of existing AI-based image synthesis methods across real-world imaging datasets spanning different modalities (MRI, mIF), phenotypes (sex, cancer subtypes, and disease status), synthesis approaches (Generative Adversarial Networks, sparse regression), and data formats (raw pixels, segmented cells). Through plasmode simulations [51] of image-phenotype associations, we demonstrate that out-of-the-box image synthesis produces invalid biological and statistical inference, observed as biased group differences. PAMS successfully mitigates these distortions in quantitative inference while remaining compatible with existing synthesis methods, bringing biologically and statistically valid synthetic imaging within practical reach.

## 2 Results

### 2.1 Synthesized images misconstrue biological findings

We used three biomedical imaging datasets to investigate the effect of using synthetic images in associations of imaging measures with participant phenotypes (Table 1).

**Table 1:** Datasets used to evaluate image synthesis. (DII: diffuse inflammatory infiltration; CLR: Crohn’s-like reaction.)

| Data Set | $N$ Patients | $N$ Images | Benchmarked Channels / Image Modality | Phenotype Context |
| --- | --- | --- | --- | --- |
| Neuroimaging | 178 | 178 | T1, Fractional Anisotropy | Sex Difference |
| Image-level mIF | 32 | 63 | CD20, CD68 | Crohn’s Disease (Active versus Inactive) |
| Single-cell mIF | 35 | 140 | CD138, CD20, CD38, CD4, CD68, CD8 | Colorectal Cancer (DII versus CLR) |

The Beijing neuroimaging open-access dataset [52] includes 178 participants (107 female) with complete fractional anisotropy (FA) data obtained from diffusion imaging. FA is a commonly used index of white matter microstructure characterizing the movement of water molecules, and higher FA values are associated with more densely packed and co-oriented white matter fibers [53]. We trained a GAN-based deep learning model [54] in a subset of 89 participants (49 female; “training data”) from the Beijing dataset to create synthetic FA images from T1-weighted anatomical images for the remaining 89 participants (58 female; “test data”)[52]. Within the test data, we compared sex differences in synthesized and real FA values. Analyses were conducted on regional mean values across 17 white matter tracts from the JHU Eve atlas white matter parcellation registered to MNI space [55]. For both the real and synthesized data, we used Welch’s t-test to perform statistical inference. The synthesized FA images were moderately similar to the true FA images when evaluated with structural similarity metric (SSIM) [56] (Figure 1a). However, FA values in major white matter tracts were qualitatively and statistically different in the synthesized images from the original: synthetic mean FA estimates had less variability across subjects compared to real images; sex differences derived from synthetic images can be qualitatively different from real images for certain regions (e.g., cingulum cingulate gyrus, Figure 1b).

**Figure 1.**
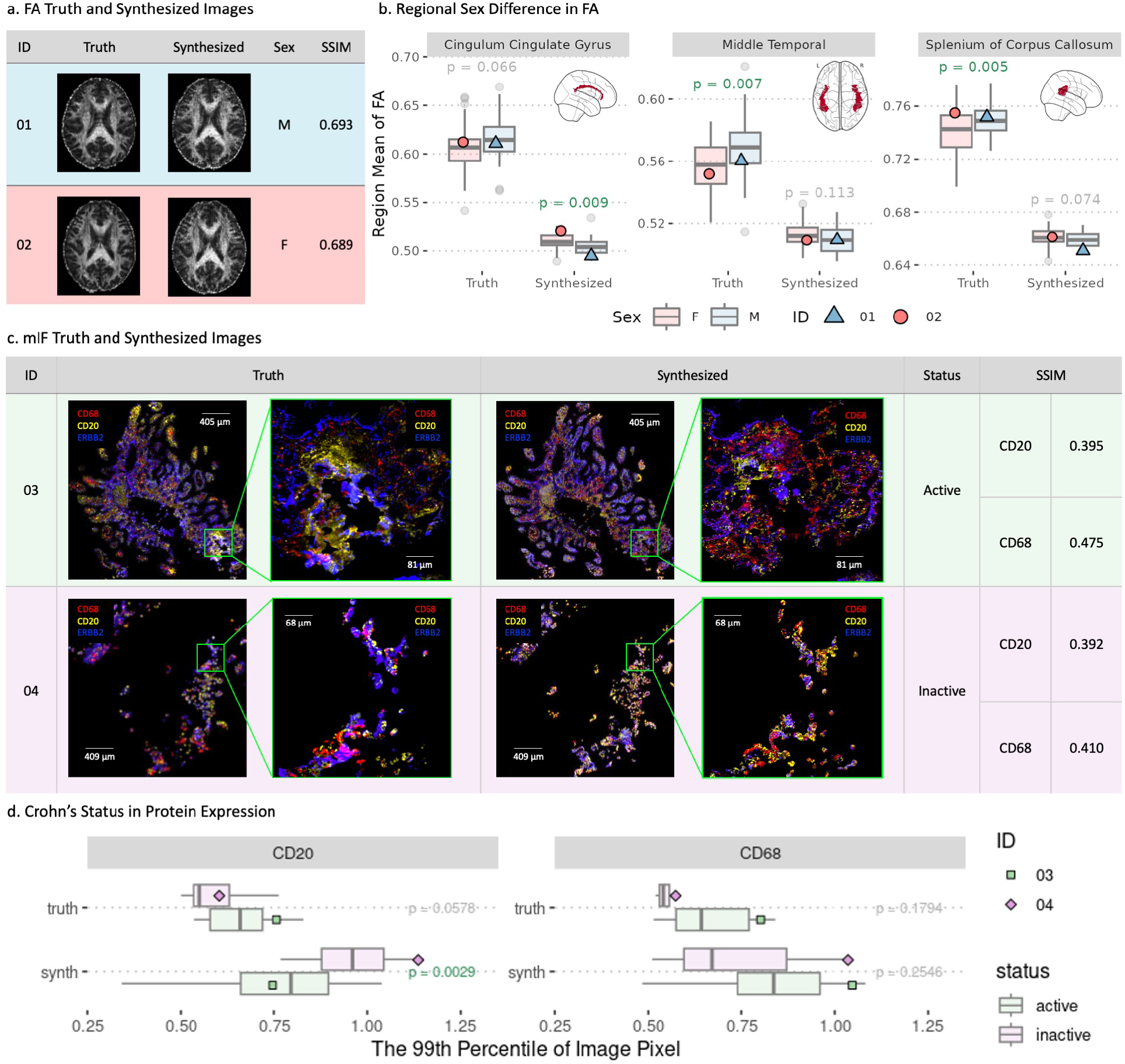
Synthetic images produce biologically misleading results compared to real data. a. Synthesized and true FA images appear visually similar in two representative patients from the Beijing dataset (M=male; F=female; SSIM=structural similarity metric). b. Synthesized FA images lead to biased estimates of sex differences across participants in regional mean FA with qualitatively different findings than the real data in major white matter tracts (green p-values are significant from t-tests). Full comparison of all regions are in Figure S1. c. Broadly, synthesized and true mIF images from two patients in a Crohn’s disease study appear similar, but there are localized differences for selected CD68 and CD20 markers shown in enlarged regions. d. The association between per-image CD68 and CD20 expressions (summarized as 99th percentile) versus active or inactive Crohn’s disease is qualitatively different based on true and synthesized images. Full comparison for all markers and different percentiles are in Figure S2.

To investigate whether the same contradictory result occurs in synthetic mIF images, we performed a similar analysis in a mIF dataset containing a panel of two immune markers for patients with Crohn’s disease (Table 1) [57]. Patients were separated into groups of active and inactive Crohn’s disease based on Crohn’s Disease Activity Index (CDAI), and patients with lower than 150 CDAI were classified as inactive [58]. The dataset included 105 tissue sections (manually isolated from full images for the ease of analysis) from 32 patients, each having 29 channels. A subset of 67 tissue sections (35 active; “training data”) were used to train an external image synthesis model pixN2N-HD [31], a multi-channel conditional GAN that uses random channel gating to synthesize missing mIF marker channels from the remaining available channels). The remaining 38 tissue sections (29 active; “test data”) were used to compare true and synthesized immune marker expression values (synthesis for each marker based on all other marker channels). We quantified differences in immune activation using B cells (CD20) [59], and macrophages (CD68)[60]. Both immune cell types have been implicated in chronic inflammatory bowel diseases such as Crohn’s disease [61, 62]. While the synthesized images appeared overall similar to real images, they failed to recapture nuanced, localized features such as the rare but important phenotypes of concentrated B cell and macrophage markers at specific tissue sections (Figure 1c). To quantitatively reflect this, we benchmarked differences in the 99th percentile of CD20/CD68 expressions between active and inactive patients on either true or synthesized images [63, 64] (99th percentile of the markers selected to reflect the rare distribution of corresponding immune cells in tissue sections). Biological conclusions were again qualitatively different between the true and synthetic images; the synthetic images had larger quantile values and the group effects were in opposite directions for real and synthesized CD20 (Figure 1d).

### 2.2 Multiple image synthesis to improve biological inference

To understand why image synthesis may cause misleading biological findings, we conceptualize it within the statistical framework of missing data imputation [49]. Valid downstream inference from synthesized images relies on two key conditions. First, it should condition on relevant covariates to ensure unbiased estimation (i.e., quantities estimated from synthesized data correctly target the biological truth), especially with respect to the phenotype of interest [65]. Bias arises when the missing images depend on, but are not synthesized using variables correlated with the phenotype, including covariates that are not directly related to the scientific question [66, 67]. For example, in the examples above, FA values are affected by sex but are imputed using only T1; missing immune marker values depend on Crohn’s disease condition but are imputed with other markers only. Such omissions thus attenuate the associations between synthesized images and the phenotypes of interest. Second, when included in downstream analyses, synthesized images should be treated as with inherent uncertainty from the synthesis procedure (as opposed to fixed, observed truth)[68]. In practice, most image syntheses methods often do not consider relevant phenotypic variables for the synthesis process or the uncertainty thereof, which can produce biased parameter and standard error estimates, leading to inaccurate biological and statistical conclusions.

As a solution, we propose PAMS, a framework that improves statistical and biological inference from synthesized images by augmenting them with considerations for key, informative covariates such as phenotypes or outcomes as well as the uncertainty during synthesis (Figure 2). PAMS introduces a stratified calibration step based on phenotype or outcome groups to recover information related to these variables (Figure 2b). Theoretically, this step ensures that the conditional distribution of the PAMS-calibrated values from synthesized images given covariates match those from the true images. The approach is fully modular: it operates on existing image synthesis output through calibration and bootstrapping of the learned error distributions that capture synthesis uncertainty, allowing seamless integration with existing image synthesis methods (see Methods).

**Figure 2.**
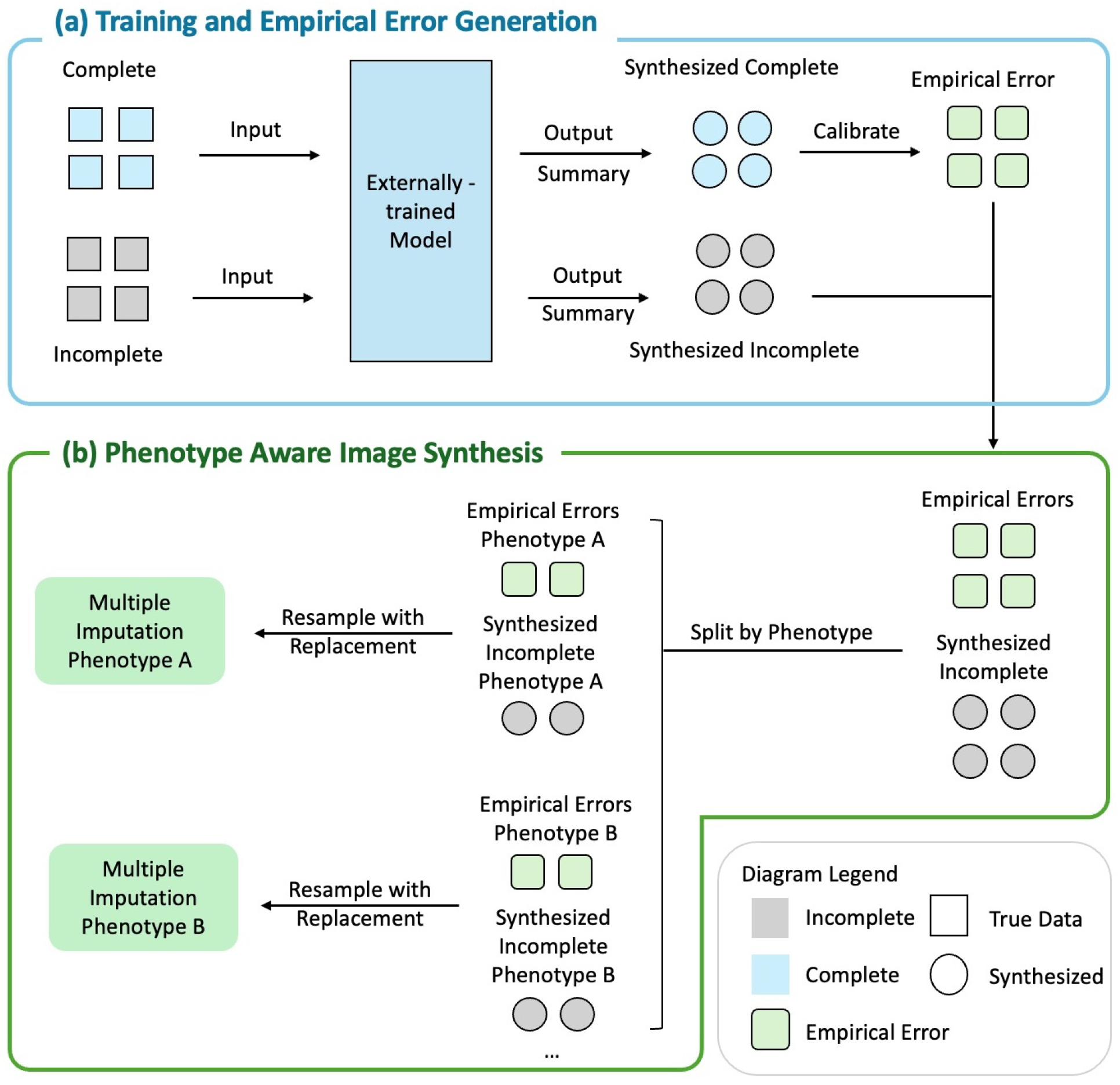
Multiple synthesis analysis pipelines. a. Complete and incomplete data are used as input to an externally-trained black-box image synthesis model. Complete data, which have true values of the synthesis model output, are used to learn an empirical error distribution (which reflects both bias and uncertainty) of the imaging values, which are in turn used to calibrate synthesized images. b. The empirical errors are stratified, bootstrapped and combined with synthesized images for multiple image synthesis (green boxes). PAMS creates multiple imaging dataset sampling errors dependent on non-imaging phenotypes, ensuring that both phenotype-dependent difference and imputation uncertainty are incorporated in downstream analyses.

### 2.3 PAMS improves biological inference of sex differences in FA

We used plasmode simulations [51] to evaluate the naive application of an image synthesis model [54] versus PAMS, which incorporates key phenotypes and imputation uncertainty to accurately inference sex differences in mean FA of subjects’ MRI images. We simulated a realistic image synthesis workflow by training a model on half of the data and evaluating them on the other half. Missing FA images were introduced randomly and synthesized, and downstream analyses were performed using robust linear regression with regional mean FA as the outcome and sex as the predictor (Figure S3). We compared three approaches: complete-case analysis (i.e., using only observed true images), image synthesis (i.e., naive analysis of synthesized images as if they were true images), and PAMS, with performance summarized across 500 bootstrapped simulations (see Methods for full details).

To evaluate the biological and statistical accuracy of the methods, we compared four metrics on their performance in estimating sex differences in mean FA (details in Methods): (1) mean sex differences in FA as estimated by the different approaches, compared against the gold standard of fully observed data results, reflecting estimation bias, (2) accuracy of model-based standard error in estimated sex differences, which reflects (mis)calibration of models’ confidence in their estimates and affects statistical inference validity (i.e., control of type-I errors, validity of confidence intervals), (3) root-mean-square error (RMSE), reflecting overall accuracy that incorporates both bias and variability, and (4) coverage probability of the 95% confidence intervals, reflecting inference validity.

Results for major white matter tracts (cingulum cingulate gyrus, middle temporal, and splenium of corpus callosum) showed that PAMS is consistently more effective at improving biological and statistical accuracy of analyses using synthesized images (Figure 3). Naive image synthesis has larger biases in estimating sex differences compared to the gold-standard, whereas complete-case analyses and PAMS had little bias. This demonstrates the importance of including phenotype information in image synthesis, especially if the downstream task is to associate images with such phenotypes; this is an issue most image synthesis approaches do not consider [69]. Correspondingly, complete-case analyses and PAMS both had best RMSEs and 95% confidence interval coverage, reflecting good accuracy in estimating sex-related FA differences (likely driven by their unbiasedness rather than reduced variability). PAMS outperformed image synthesis, which had significant undercoverage, again likely driven by their severe bias in estimating true sex differences. Overall, our findings indicate that PAMS’s modular incorporation of phenotype information in image synthesis ensures valid downstream statistical and biological findings. We extended the analysis to the other 15 white matter regions (Figure S4) and we found that PAMS performs consistently across the other regions.

**Figure 3.**
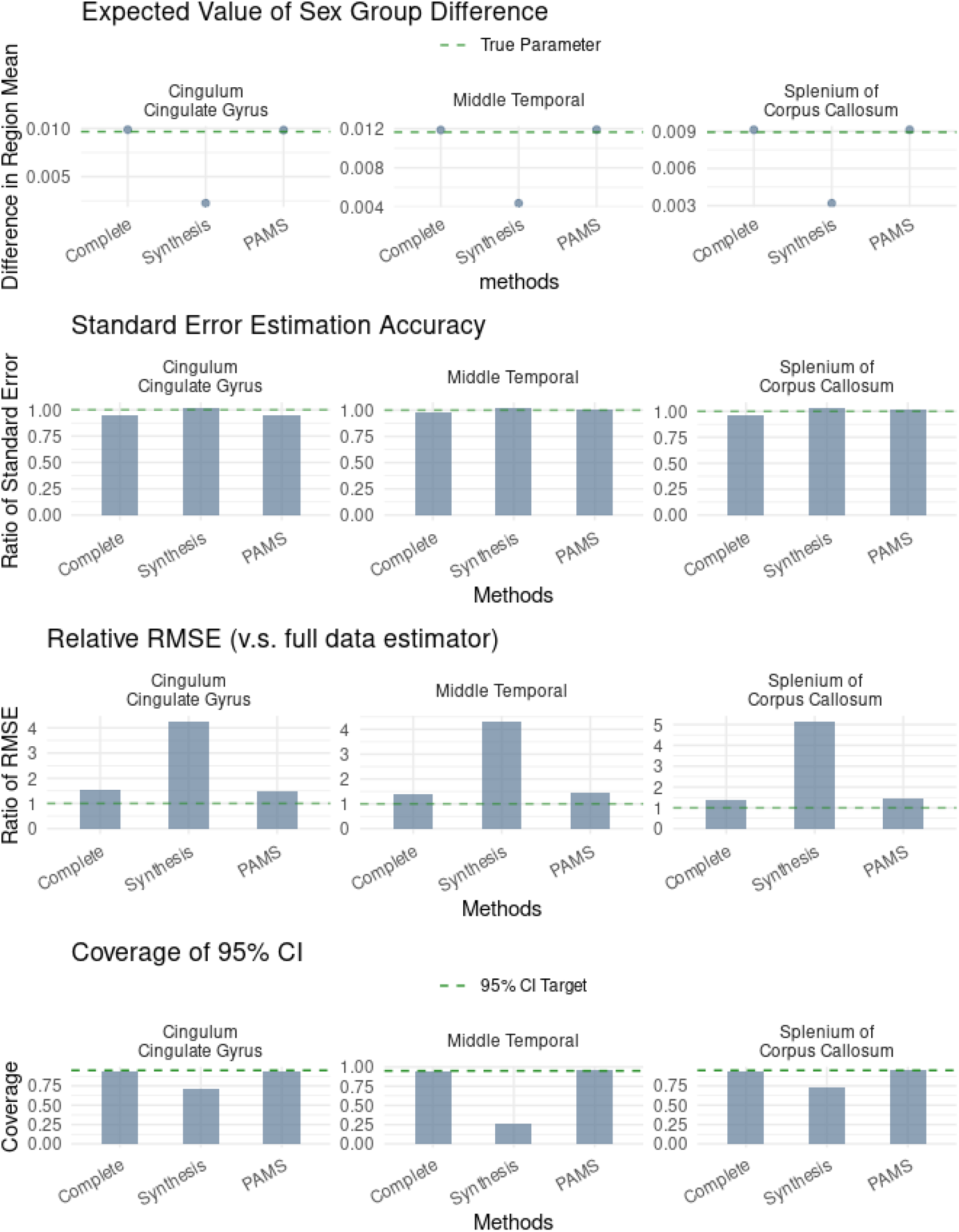
Bootstrap analysis of sex differences in fractional anisotropy (FA) in major regional white matter regions. Evaluated approaches include (1) analyses using only observed true images (i.e., complete case analysis, “Complete”, (2) naively incorporating images synthesized from a GAN-based synthesis approach as if they were true, observed images (“Synthesis”), and (3) our Phenotype-Aware Multiple Synthesis (“PAMS”) which augments the GAN-synthesized model. a. Only complete cases and PAMS unbiasedly estimate the true sex differences. b. All synthesis methods accurately estimated the standard error of the sex difference. c. PAMS has similar RMSE to complete case analysis indicating it is unbiased for the true sex difference and has similar variability to the baseline complete case analysis. d. Overall, PAMS confidence interval coverage is near the 95% target and comparable to complete case analysis. Image synthesis fails to reasonably capture the true sex differences in FA.

### 2.4 PAMS improves biological inference of Crohn’s disease

To validate the performance of our proposed multiple image synthesis approaches in other biomedical imaging applications and modalities, we adopted the same simulation pipeline and evaluation metrics to analyze the mIF dataset (Methods). We used a robust linear regression model for estimation of marker expression level differences between active and inactive subjects and evaluated differential expression of CD68 and CD20 in active versus inactive Crohn’s disease patients (Figure 4). Similar to the neuroimaging dataset, PAMS performed well in estimation accuracy while image synthesis produced biased results (Figure 4a). PAMS has a standard error compared to complete case analysis (Figure 4b), which leads to a relatively higher RMSE (Figure 4c). The coverage of PAMS is above 95% and is slightly higher than complete case analysis (Figure 4d). Coverage using image synthesis is very low for CD20, caused by the magnitude of the bias. These trends were consistent for other key markers (Figure S5).

**Figure 4.**
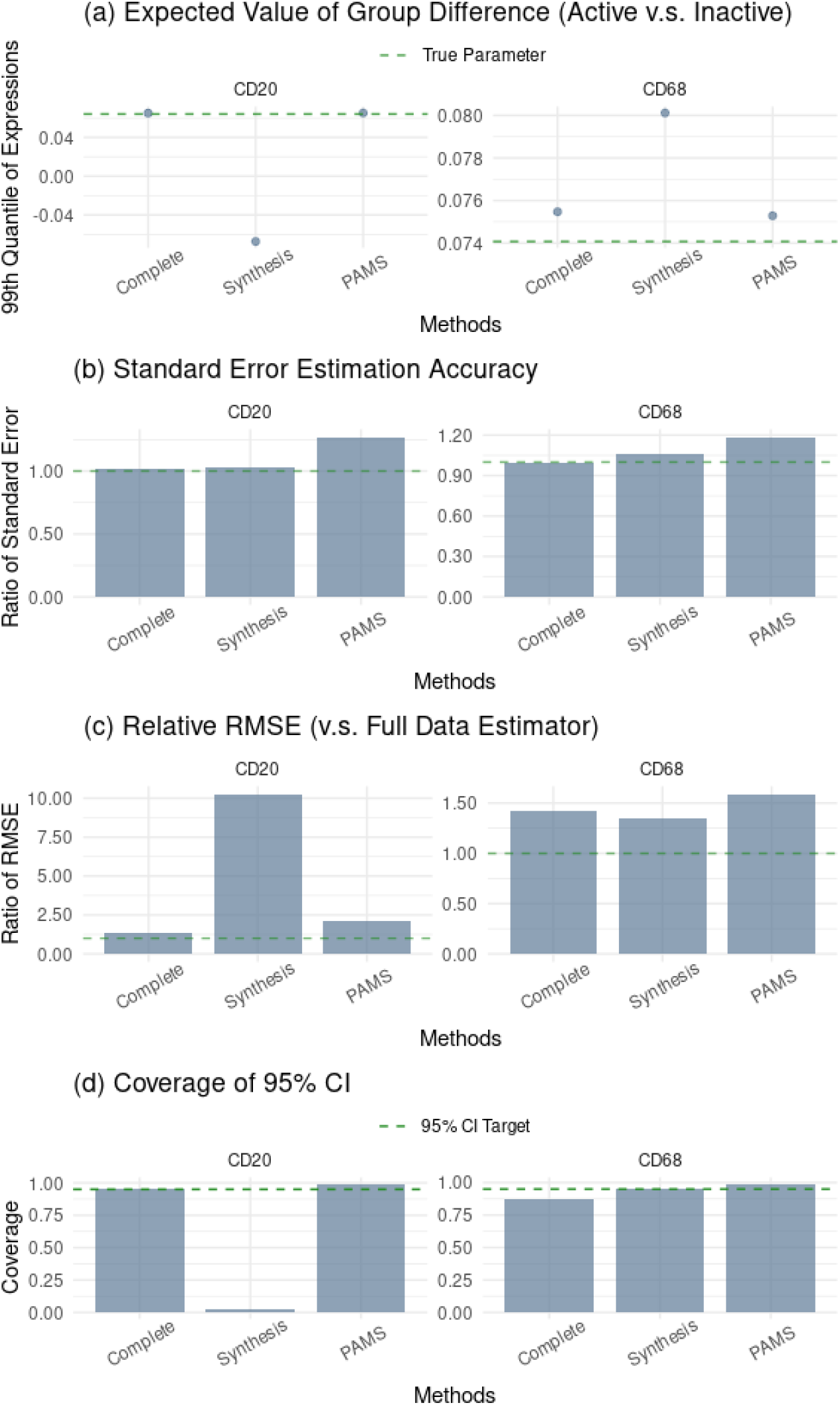
mIF image-level bootstrap analysis of disease status differences (active versus inactive) in the 99th percentile expression comparing complete case analysis (“Complete”) to the synthesis analysis methods (GAN-based naive synthesis by pixN2N-HD [31], “Synthesis”; PAMS augmentation, “PAMS”). a. Only complete case analysis and PAMS unbiasedly estimate the true disease difference for both markers. b. PAMS slightly overestimates standard error, but c. has much smaller RMSE relative to naive synthesis for CD20 and slightly higher RMSE than complete case analysis for both markers. d. Only PAMS consistently has above-nominal coverage.

### 2.5 PAMS improves biological inference of cancer prognosis

We also explored how the multiple synthesis principle applies to a mIF tissue microarray (TMA), where the tissue area is from smaller tumor-only tissue cores and applied to processed cell-level data (Methods). This study examined 140 TMAs for advanced-stage colorectal cancer (CRC) patients with different tumor types: “Crohn’s-like reaction” (CLR) and diffuse inflammatory infiltration (DII) (Table 1) [70]. The study found that immune markers for cytotoxic T cells (CD8) and macrophages/monocytes (CD68) had different expression levels between these two tumor types. We trained a Lasso model on 70 samples to impute per-cell missing channels from observed channels and used the remaining 70 samples for evaluation (see Methods). Biological analyses compared cell-level immune marker expression level as the outcome, with tumor type as the independent variable in robust linear regression models. Evaluated markers include CD138, CD20, CD38, CD4, CD68, CD8, which are most closely related to the tumor types in the original study [71]. The methods were evaluated here using the same criteria established in the preceding sections. Again, PAMS was less biased compared to image synthesis (Figure 5a). PAMS slightly underestimated standard errors compared to complete case analysis and image synthesis (Figure 5b), reflecting not only improved bias compared to image synthesis, but also efficiency gain (i.e., smaller variance) by including synthesized images in addition to real data. Image synthesis had inconsistent RMSE, with the best efficiency for CD38, CD4 and CD8, and the worst for CD138, CD20 and CD68 (Figure 5c). However, PAMS consistently has better RMSE than complete case analysis and the most consistent confidence interval coverage across the markers, despite slightly underestimating standard errors (Figure 5d). The observed trends extended to other key markers. (Figure S6)

**Figure 5.**
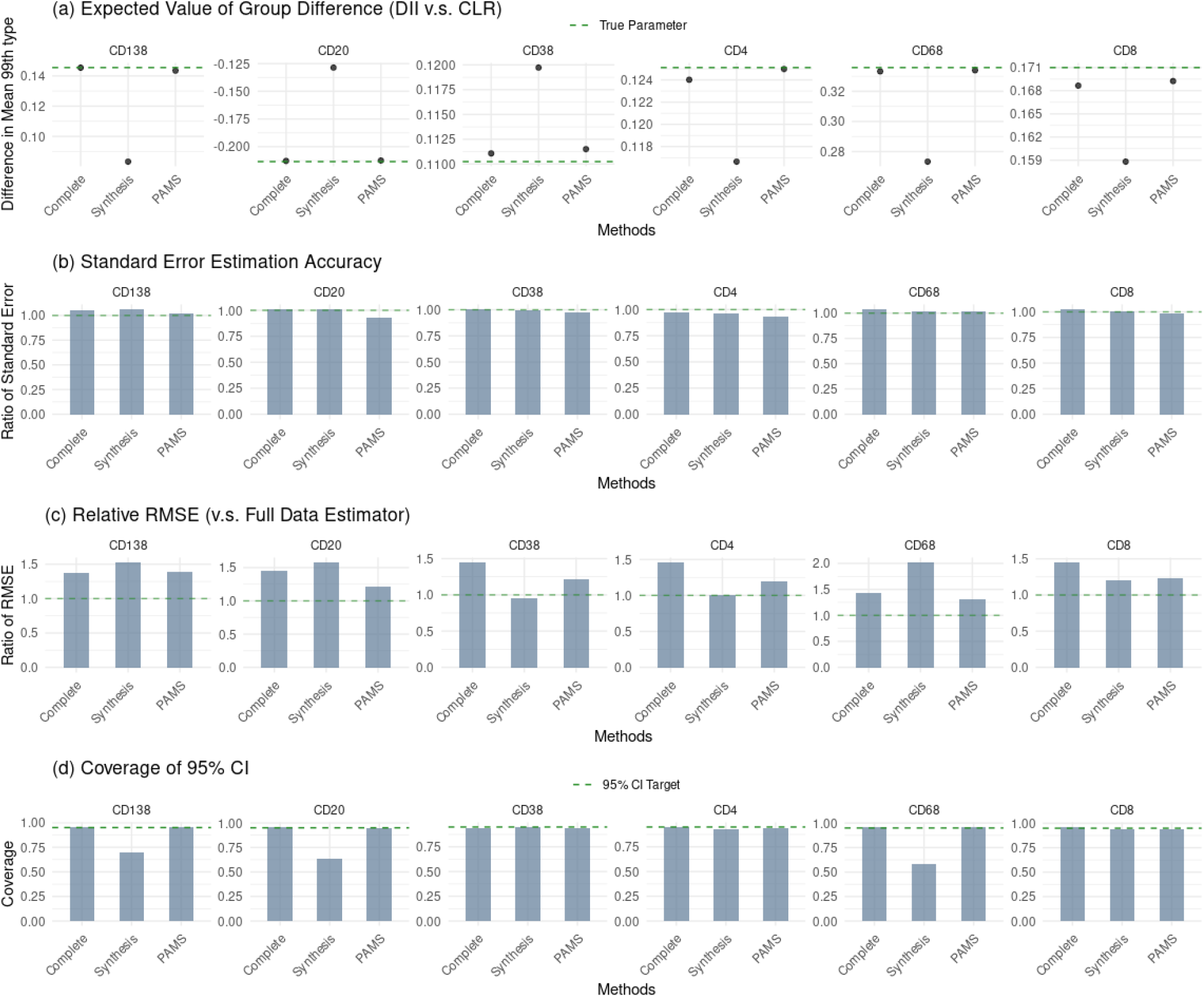
mIF cell-level bootstrap analysis of cancer type differences (CLR vs. DII) in the 99th percentile expression comparing complete case analysis (“Complete”) to the synthesis analysis methods (Lasso-based synthesis, “Synthesis”; PAMS). **a.**Complete case analysis and PAMS are much less biased in estimating the true disease difference for all markers compared to synthesis analysis. b. PAMS slightly underestimates standard error, but c. has consistent lower or comparable RMSE than complete case analysis for all markers. For CD38, CD4 and CD8, multiple synthesis has a lower RMSE than the synthesis analysis methods. d. Both PAMS and complete case analysis consistently have above-nominal coverage.

## 3 Discussion

We evaluated the biological and statistical validity of analyzing synthetic images, showing that failing to incorporate phenotype in the synthesis algorithm and ignoring uncertainty inherent to the synthesis methods can introduce statistical bias and lead to invalid biological conclusions. Although image synthesis is widely studied and algorithms are being made available [21, 25], few studies have investigated the validity of downstream analyses based on these synthesized images, despite recognition of its importance [37]. Most synthesis methods do not account for phenotype or outcome variables during the synthesis step. Drawing from well-established missing data statistical theory, unless image synthesis is near-perfect, omitting outcome information during imputation will yield biased findings, a pattern we consistently observed across almost all cases when synthesized images were naively incorporated.

To address this, we introduced PAMS, a modular solution that corrects such biases through estimating the error process conditioning on the phenotype/outcome of interest, without requiring modification to the underlying synthesis method. At the same time, PAMS estimates the error process to appropriately capture the uncertainty in the synthesized image. Together, these contributions represent one of the first efforts and a proof-of-concept for combining existing image synthesis approaches with established missing data statistical theory, a practice that likely will be crucial as the field moves beyond image synthesis method development to deployment in scientific research.

Across our evaluated application cases, PAMS consistently addressed the bias induced by image synthesis. However, in two cases (sex difference MRI imaging, Crohn’s disease phenotypes associated with pixel-level mIF), it did not lead to performance improvement (i.e., smaller RMSE) compared to complete case analyses (i.e., analyses using only observed images). Only in analyzing CRC phenotypes as associated with cell-level mIF expressions did we observe a consistent performance gain. This likely indicates the low-quality of the synthesized images in targeting the quantities of interest in the two cases (average FA values; tail percentile of pixels’ immune marker values), as noted via direct inspections (1). Our findings here echo previous concerns that, while synthesized images visually and computationally appear similar to real ones, they can still perform poorly in approximating key biological characteristics of their targets [37]. Under such regimes, PAMS cannot improve beyond complete case analyses given the poor quality of synthesized images. Nevertheless, it can still safeguard the bias they might cause when naively included in the analysis.

While PAMS is among the first to address biological validity in the context of synthesized biomedical images, a small body of work has acknowledged similar issues and proposed solutions in various application areas. For instance, prediction-powered inference and conformal prediction both considered the variance from the machine learning process, aiming to either quantify variability or create prediction intervals. Notably, Sun et al [46] incorporated variance pooling from the multiple imputation framework for the downstream analysis of spatial gene expression prediction, reflecting a related awareness of synthesis uncertainty. However, these studies only focused on one of bias and variation in the synthesis process, while PAMS reduces both bias and underestimation of synthesis variation through a modular framework. The key novelty of PAMS is brought by stratified sampling of error conditioning on phenotype, which allows arbitrary synthesis models while also correcting biases.

PAMS offers one solution to ensuring the biological utility of synthetic images, though its application comes with important considerations. First, PAMS cannot fully compensate for limited synthesis accuracy with respect to the downstream quantity of interest. Although this does not cause bias in PAMS-based findings given its design, it will lead to efficiency loss, as observed in our analysis with image-level mIF marker CD20. Conversely, high synthesis accuracy alone does not eliminate the need to consider the phenotype, model error, and error process when synthesized images are used for scientific research. Second, our current solution assumes that the distribution of prediction errors are identical across image expression values. This assumption may not hold for all datasets, especially for irregular shaped tissues such as mIF imaging, where spatial distribution of cells varies greatly with tissue size and shape. Despite these considerations, our work highlights the importance of evaluating image synthesis methods beyond the typical protocol of assessing visual and quantitative accuracy, emphasizing that biological and statistical validity are critical prerequisites for image synthesis to have utility in scientific research.

## 4 Methods

### 4.1 Data

The neuroimaging data were obtained from the publicly available Beijing Enhanced DTI dataset hosted on NITRC [52, 72, 73]. The dataset comprises diffusion tensor imaging (DTI) and structural T1-weighted MRI scans from healthy participants, collected as part of an open neuroimaging initiative for population-level brain studies. Each subject contributes a high-resolution T1 image and a corresponding DTI scan with precomputed fractional anisotropy (FA) maps; the dataset is widely used for benchmarking owing to its standardized acquisition and open accessibility [74, 75]. T1 images were skull-stripped and nonlinearly registered to the MNI152 1 mm template using FSL and C3D [76, 77], and FA images were aligned to the same template. Both modalities were resliced from 1 mm^3^ to 2 mm^3^ isotropic voxels.

Two multiplex immunofluorescence (mIF) datasets were used to assess generalizability. The *pixel-level mIF* dataset [57] are whole-slide samples from terminal ileum (TI) and ascending colon (AC) tissues in patients with active and inactive Crohn’s disease. Tissue imaging is performed cyclically, where the tissues were stained with marker-specific fluorescent antibody dyes, scanned and photo-inactivated. The images are then computationally processed for registration and illumination correction.The *cell-level mIF* dataset is publicly available through Mendeley Data [71] and consists of tissue microarray samples from human colorectal cancer, stratified by the presence or absence of tertiary lymphoid structures (TLS). The cell-level data were processed as previously described [70]: light deconvolution, background subtraction, manual cell segmentation, and marker gating. For both mIF datasets, fluorescence intensities were normalized by slide-specific mean division followed by log_10_ transformation to reduce between-slide and between-batch variability [78].

### 4.2 Missing-data framework for image synthesis

We formalize statistical inference with synthesized images using the missing data framework [49, 50]: the full analytical dataset includes all subjects, with the underlying images unobserved for the subset requiring synthesis. For analytic simplicity, we initially assume image missingness operates under missing completely at random (MCAR) [47]: the missingness of an image is independent of all observed and unobserved variables, including the underlying image trait *Y* \, the phenotype *X*, the synthesizer’s input *W* \, and additional covariates *Z*. MCAR is a strong assumption and is unlikely to hold exactly in practice — for example, in cyclic mIF staining, images may be lost to tissue degradation in batches that align with phenotype groups, producing dependence between missingness and *X*. We adopt MCAR here for clarity of exposition and relax it to missing at random (MAR) in Section 4.2.6.

### 4.2.1 Notation

Let *Y* denote the image-derived quantitative trait of interest (e.g., a regional mean FA, or a percentile of pixel intensities across an mIF image). Let *Y ^*^* denote the corresponding trait computed from a *synthesized* image. Synthesized images are produced by an externally trained synthesizer *f* that takes as input an auxiliary modality or data source *W* (e.g., the T1 image when synthesizing FA, or the remaining marker channels when synthesizing a held-out channel), so that *Y ^*^* = *f* (*W*) for a learned *f* . We assume *W* is observed for all subjects. The learning for *f* is usually enabled through existing algorithms on external data; our method is proposed as a modular improvment given the learned *f* and synthesized *Y ^*^*. The specific choices for *f* in our application cases are detailed in Section 4.3.1.

Let *X* denote the phenotype of interest (e.g., sex, disease status) and *Z* denote additional covariates requiring adjustment (e.g., age, BMI). Although the framework is stated at the image level for clarity, it extends to pixel- or cell-level synthesis. Without loss of generality, we order the *n* subjects so that the first *n*^obs^ have observed images and the remaining *n*^synth^ = *n* − *n*^obs^ have missing images requiring synthesis.

Thus *Y*_*i*_ is observed for *i* = 1, …, *n*^obs^ and missing for *i* = *n*^obs^ + 1, …, *n*; the corresponding synthesized values 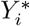are available for all *i* but are used only in place of missing *Y*_*i*_. The phenotype *X*_*i*_ and covariates *Z*_*i*_ are observed for all *i* (we consider only image missingness).

The target of inference is a parameter *θ* relating *Y* to *X*, potentially adjusted for *Z*. For all our applications, *θ* is specifically a *mean difference* parameter. We give the unadjusted form below for each application; each can be extended to a covariate-adjusted contrast by conditioning on *Z* (e.g., *θ* = E[*Y* | *X* = 1, *Z*] −E[*Y* | *X* = 0, *Z*] averaged over *Z*, or fit via a regression of *Y* on *X* adjusting for *Z*):

1. *Neuroimaging. X* ∈ {0, 1} indicates sex and *Y* is a subject’s mean FA value within a specified white matter tract; the target is *θ* = E[*Y* | *X* = 1] − E[*Y* | *X* = 0].
2. *Pixel-level mIF. X* ∈ {0, 1} indicates Crohn’s disease activity (1 = active, 0 = inactive control) and *Y* is the 99th percentile of an immune-marker channel; the target is *θ* = E[*Y* | *X* = 1] − E[*Y* | *X* = 0].
3. *Cell-level mIF. X* ∈ {0, 1} indicates TLS-related tissue context (1 = CLR, 0 = DII) and *Y* is the 99th percentile of an immune-marker channel; the target is *θ* = E[*Y* | *X* = 1] − E[*Y* | *X* = 0].

### 4.2.2 Complete-case estimator

If all data were observed, the typical procedure is to apply some standard estimator *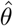* to all data {(*Y*_1_, *X*_1_, *Z*_1_), …, (*Y*_*n*_, *X*_*n*_, *Z*_*n*_)} to estimate *θ*. For example, in our mean difference applications,

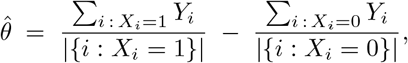

which is the difference of subgroup means in the full sample.

We further restrict our discussion to estimators *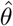* that are *asymptotically linear*. That is, *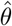* can be written as

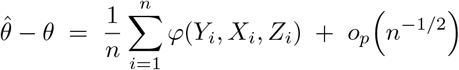

for some influence function *φ* satisfying E[*φ*(*Y, X, Z*)] = 0. By the law of large numbers, this representation implies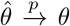. Asymptotically linear estimators cover many typical estimation scenarios, including mean differences, (generalized) linear regression coefficients, and Cox model log-hazard ratios. For example, for the mean difference above, the influence function is

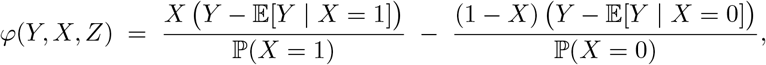

which in this unadjusted case does not depend on *Z*.

In the presence of missing data,*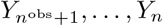* are unobserved. A natural first approach is the so-called *complete-case estimator*, which applies *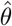* only to the subjects with observed images:

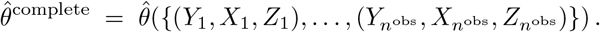

By asymptotic linearity, it is easy to see why *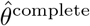* is valid for *θ*. Under MCAR, the labelling of subjects as missing vs. observed is independent of (*Y, X, Z*), so {(*Y \, X \, Z*)} 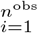 is itself an i.i.d. sample from the population distribution of (*Y, X, Z*). Asymptotic linearity therefore applies directly with sample size *n*^obs^:

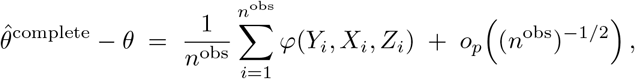

and the law of large numbers gives *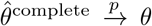* . The deficit of complete-case relative to the full-data estimator is therefore not bias but *efficiency* : it discards the *n*^synth^ subjects with synthesized images, which would otherwise contribute to a larger effective sample size and lower estimation variance. The PAMS estimator developed in Section 4.2.4 aims to improve this inefficiency without sacrificing consistency.

Specifically for the mean difference example, the complete-case estimator reduces to:

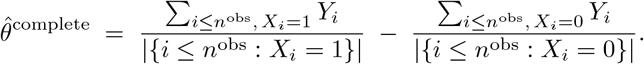

### 4.2.3 Naive synthesis estimator

A natural alternative to discarding subjects is to fill missing images with their synthesized counterparts and apply *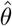* to the resulting pooled dataset. We call this the *naive synthesis estimator* :

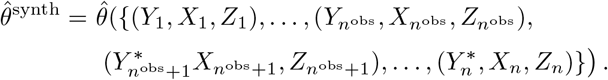

By asymptotic linearity of *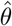* applied to the pooled sample,

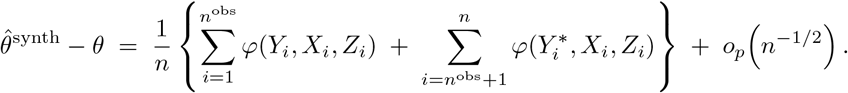

Under MCAR the first sum is mean-zero by the argument above. The second sum, however, evaluates *φ* at the synthesized value *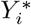* in place of the true *Y*_*i*_. Because the joint distribution of (*Y ^*^, X, Z*) generally differs from that of (*Y, X, Z*), or, more specifically, the conditional distribution *Y ^*^*|(*X, Z*) typically differs from that of *Y* |(*X, Z*), we have that

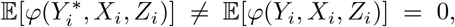

and *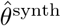* is generally inconsistent for *θ*.

Specifically, for the mean difference case,

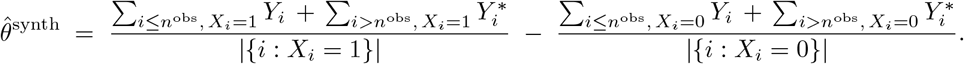

The bias has a clean and interpretable form: substituting the influence function and taking expectations gives

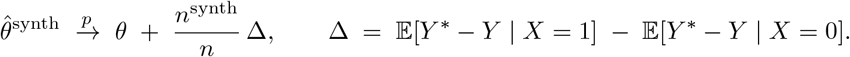

The bias scales with two quantities: the fraction of synthesized observations *n*^synth^*/n*, and the *differential synthesizer bias* ∆ — the difference between the synthesizer’s mean error within the two phenotype groups. As we demonstrate below, the PAMS estimator developed next is designed to eliminate the ∆ bias.

### 4.2.4 PAMS estimator

The above sections illustrated the trade-off between the two simple estimators: 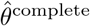is consistent but disregards all potentially useful synthesized images *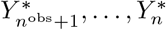*, while *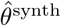* uses these synthesized images but in a naive fashion that introduces bias, as 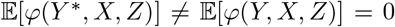. Our proposed PAMS estimator *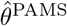* resolves this trade-off by post hoc correcting each *Y ^*^* into an imputed value, *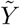* \, whose conditional distribution given (*X, Z*) matches that of *Y* \, so that 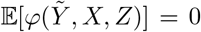and the bias vanishes while the synthesized subjects still contribute to estimation. Briefly, we construct empirical residuals *Y* − *Y ^*^* on the observed-image subjects, draw them stratified by (*X, Z*) and add them back to *Y ^*^*, embedded in a multiple imputation framework to also account for imputation variability. We formally establish the procedure to construct *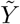* \, the corresponding PAMS estimator *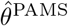*, and its validity below.

**Generating** *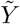* For *i* = 1*, …, n*^obs^, define the empirical synthesizer residual

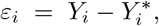

which is computable since both *Y*_*i*_ and *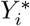* are available for observed-image subjects. We then generate *M* imputed datasets indexed by *m* = 1*, …, M* . For each *m* and each subject *i* = *n*^obs^ + 1*, …, n* with missing image, we draw an empirical residual *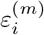*from a non-parametric estimate of the conditional residual distribution *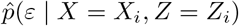*, and form the imputed value

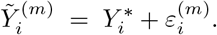

The form of*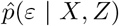*depends on the structure of the conditioning variables. When the conditioning set consists only of the phenotype *X* — as in our three applications, where the analysis is an unadjusted mean difference between two groups — 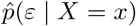 is the empirical distribution within the *X* = *x* stratum for *x* ∈ {0, 1}, and the residual draw reduces to a phenotype-stratified bootstrap:

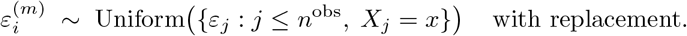

The same construction extends to conditioning on (*X, Z*). For categorical *Z*, the stratified bootstrap generalizes directly with strata defined by (*X, Z*) cells.

**Multiple imputation estimator** *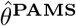* For each imputation *m*, we compute *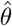* on the pooled dataset with *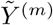*in place of the missing *Y* :

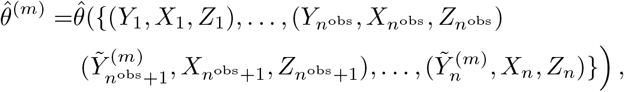

together with its standard error SE(*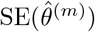*). The PAMS point estimate is the average across imputations,

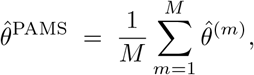

and its standard error is obtained by Rubin’s rules [47, 79]. With within-imputation variance

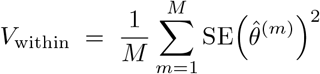

and between-imputation variance

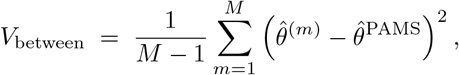

the pooled standard error is

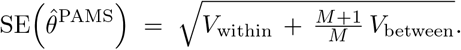

**Consistency of** *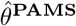* For intuition, consider the single-imputation version of PAMS, in which one draw *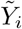*replaces each missing *Y*_*i*_ before estimation. Under MCAR, by asymptotic linearity of *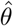* applied to the pooled sample,

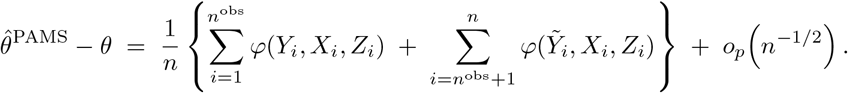

The first sum is mean-zero by the complete-case argument of Section 4.2.2. For the second sum, *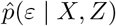* converges to the true conditional residual distribution as *n*^obs^ → ∞, so *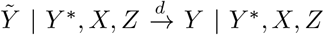*. By the law of iterated expectations,

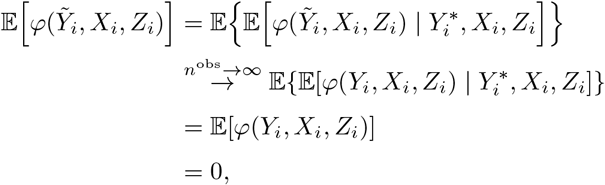

and consequently *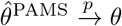 θ*.

If the synthesizer is strong enough that *Y* ⊥ (*X, Z*) | *Y ^*^* holds — i.e., *Y ^*^* captures all of the variation in *Y* that is associated with (*X, Z*) — then stratification is unnecessary and an unstratified empirical residual bootstrap is also consistent. This regime is the basis for the multiple synthesis comparator in Section 4.2.5.

#### 4.2.5 Multiple synthesis estimator

We additionally evaluate a *multiple synthesis* estimator, *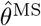*, as a comparator that accounts for synthesis variability without phenotype stratification. Multiple synthesis follows the same multiple imputation framework as PAMS (Sections 4.2.4–4.2.4) but draws each residual from the *marginal* (unstratified) empirical distribution 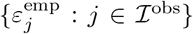. By the argument at the end of Section 4.3.3, *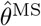* is consistent under the stronger assumption *Y* ⊥ *X* | *Y ^*^*; PAMS relaxes that assumption.

#### 4.2.6 Relaxing the MCAR assumption to MAR

The MCAR assumption simplified the derivations above but is unrealistic in practice. We now extend the framework to missing at random (MAR), under which the missingness of an image may depend on observed variables but not on the unobserved image itself:

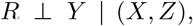

where *R* ∈ {0, 1} indicates whether the image is observed. MAR is strictly weaker than MCAR and is the standard working assumption in missing data analysis [47]. We distinguish two regimes based on whether the analyst’s estimator targets a functional of the conditional distribution *P* (*Y* | *X, Z*).

##### Targets that are functionals of

*P* (*Y* | *X, Z*) When the target *θ* is a functional of the conditional distribution *P* (*Y* | *X, Z*) alone — equivalently, when the influence function satisfies the conditional mean-zero property

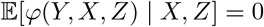

— the MCAR consistency arguments carry over to MAR without modification. This class is the standard one for which complete-case analysis remains valid under MAR [47, 50]; it covers broad estimation procedures in practice, including e.g. unadjusted and covariate-adjusted mean differences (all three applications in this paper), (generalized) linear regression coefficients under correct specification of the conditional mean, and Cox model log-hazard ratios under correct specification of the conditional hazard.

##### Complete-case estimator

The asymptotic-linearity expansion of Section 4.2.2 gives

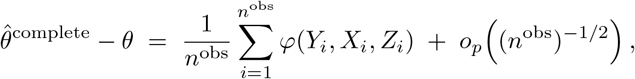

which converges in probability to E[*φ*(*Y, X, Z*) | *R* = 1]. By the law of iterated expectations and MAR,

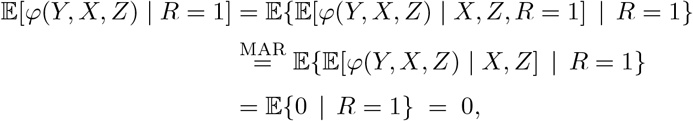

where the second equality drops *R* from the inner conditioning by MAR (*Y* ⊥ *R* | *X, Z*), and the third uses the conditional mean-zero property of *φ*. Hence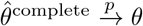.

##### PAMS estimator

The PAMS expansion of Section 4.2.4 contains two sums. The observed-subject sum is mean-zero by the complete-case argument above. For the synthesized-subject sum, the empirical residual distribution *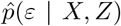* is constructed from observed-image subjects but used to impute missing-image subjects, so its validity rests on the residual distribution agreeing between the two subsamples. Under MAR, *Y* ⊥ *R* | (*X, Z*) implies that *Y* − *Y ^*^* has the same conditional distribution given (*X, Z*) regardless of *R*, so *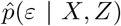* targets the correct conditional residual distribution for the missing-image subjects. The convergence *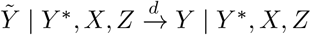*therefore holds as under MCAR, and the tower-property argument gives *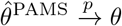*.

##### Targets that are functionals of the joint distribution

When the target *θ* is a functional of the joint distribution of (*Y, X, Z*) rather than of *P* (*Y* | *X, Z*) alone (for instance, population-marginal contrasts under non-collapsibility such as marginal odds ratios from logistic regression), the conditional mean-zero property fails. Under MAR, the observation indicator *R* shifts the distribution of (*X, Z*) in the observed subsample, so E[*φ*(*Y, X, Z*) | *R* = 1] no longer collapses to E[*φ*(*Y, X, Z*)] = 0, and *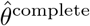* is biased.

This can be addressed straightforwardly with inverse probability weighting[80, 81] : estimate the per-subject observation probability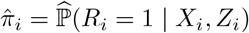 on the full sample, and weight each observed subject’s contribution to *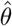* by *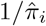* The IPW identity

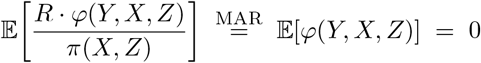

restores marginal mean-zero of the weighted influence function and hence consistency, regardless of whether the conditional mean-zero property holds. The same weighting can be applied within each PAMS imputation, replacing the unweighted *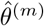* in Section 4.2.4 with its IPW analogue. We do not pursue IPW extensions empirically in this paper, since all three applications fall in the conditional-target regime, but the framework above accommodates them with no structural change.

### 4.3 Plasmode simulation

Each dataset was partitioned into a training subset, used to fit an external image synthesis model, and a simulation subset, used for plasmode evaluation [51]; the partitions are all performed at the patient level. The per-phenotype splits are given in Table 2, where each cell of Table 2 reports the number of independent subjects.

**Table 2:** Per-phenotype split of training and simulation subjects for each dataset.

| Dataset | Phenotype | Training | Simulation |
| --- | --- | --- | --- |
| Neuroimaging | Male | 40 | 31 |
|  | Female | 49 | 58 |
|  | Total | 89 | 89 |
| Cell-level mIF | CLR | 37 | 31 |
|  | DII | 33 | 39 |
|  | Total | 70 | 70 |
| Pixel-level mIF | Active | 13 | 7 |
|  | Inactive | 9 | 3 |
|  | Total | 22 | 10 |

For each dataset, we first computed the target parameter *θ* on the complete simulation subset (all images observed) to establish a per-dataset ground truth. We then generated *B* = 500 plasmode replicates per dataset. Within each replicate, we sampled 200 subjects with replacement from the simulation subset and induced missingness by randomly removing the imaging data of 50% of those subjects. The trained synthesis model produced 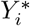 for each subject whose image was removed; residuals 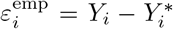 were computed on the remaining (non-missing) subjects within the replicate to populate the empirical residual distribution required by *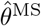* and *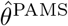* (Figure 2a). Multiple imputation was then carried out with *M* = 500 imputations for both estimators (Figure 2b); the corresponding multiple-imputation point estimates and standard errors are pooled by Rubin’s rules as in Section 4.2.4.

#### 4.3.1 Image synthesis methods

FA images in the neuroimaging dataset were synthesized using the method proposed in Gu et al [54]. This paper adapted CycleGAN [82], a deep learning method widely used for 2D image translation, to translate images from structural space (T1) to diffusion space (FA). Here, CycleGAN is extended to the 3D neuroimaging by synthesizing 2D slice-wise. That is, the model is trained with a batch of slices from the FA and T1 pairs of a certain axial coordinate, and the model synthesizes the corresponding FA slice with the same axial coordinate.

Image synthesis for pixel-level mIF dataset was carried out using the deep synthethic learning framework introduced in Bao et al [31] named pixN2N-HD. Based on pix2pixHD, a conditional GAN for high-resolution image-to-image translation, pixN2N-HD extends the framework to multi-channel synthesis using a random-gating strategy, enabling a single model to handle different missing-marker configurations. The images were split into 1024 × 1024 patches without resampling, and patches with less than 5% non-zero intensity pixels were removed.

We synthesized images in the cell-level mIF dataset with reference to the procedure outlined in Wu et al[30], where biomarker values are generated with XGBoost [83]. Despite Wu’s adoption of XGBoost with their dataset, we discovered in our preliminary work that LASSO regression [84] renders better prediction accuracy and is less computationally expensive than XGBoost. Therefore we used LASSO to synthesize the cell expression intensities in the cell-level mIF dataset.

### 4.4 Statistical analysis

Within each plasmode replicate *b* ∈ {1*, …, B*} we fit a robust linear regression matched to the original scientific question for each dataset (Table 1):

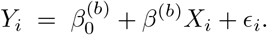

For the neuroimaging dataset, *Y* is the regional mean FA and *X* is sex. For the cell-level mIF dataset, *Y* is the 99th percentile of tissue pixel expression and *X* is the CLR vs. DII contrast. For the pixel-level mIF dataset, *Y* is the 99th percentile of pixel expression and *X* is Crohn’s disease status (active vs. inactive). For each replicate we record the point estimate *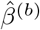* and its standard error SE*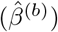*Estimates for the multiple synthesis and PAMS estimators are obtained by the multiple-imputation procedure of Section 4.2.4; the complete-case estimator *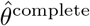* is computed on the observed-image subset only.

### 4.5 Evaluation metrics

For each estimator we report four metrics aggregated across the *B* replicates.

1. Mean point estimate.

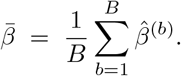
2. Standard error accuracy.

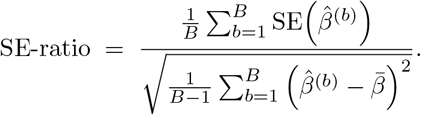

A ratio above 1 indicates conservative standard errors; below 1, anti-conservative.
3. Relative root mean squared error. Let *θ*_*b*_ denote the regression coefficient fit on the complete (all-images-observed) version of plasmode replicate *b*. Then

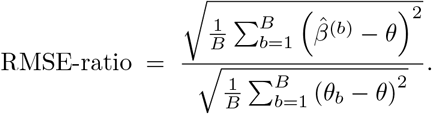

The numerator is the RMSE of the estimator against the overall target *θ*; the denominator captures irreducible bootstrap-induced variability and acts as a floor.
4. Coverage of the nominal 95% confidence interval. The proportion of replicates in which the 95% confidence interval for *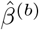* covers *θ*. Confidence intervals are constructed from the multiple-imputation standard error (Section 4.2.4) with the Barnard–Rubin small-sample degrees-of-freedom correction [85].

## 5 Funding Acknowledgments

This research was supported by NIH grants R01MH123563 (to S.N.V.), R35GM162668 (to S.M.), R01DK103831 (to K.S.L.). This research was partially supported by the Vanderbilt Department of Biostatistics Faculty Pilot Grant.

## 6 Author Contributions

S.V. and S.M. conceived the study. J. X. led data analysis, simulation studies, figure creation, and manuscript drafting. J.X., S.V. and S.M. wrote the manuscript. K.S.L, L.A.C, K.T.W collected the mIF pixel-level imaging data. S.B. and B.A.L synthesized the mIF pixel-level images. Q.L. provided technical expertise in Crohn’s disease biomarker analysis. S.V. and S.M. supervised the project, and revised the manuscript. K.S.L, S.V. and S.M. secured funding. All authors read and approved the final manuscript.

## 7 Data Availability

The neuroimaging data is publicly available at NITRC https://www.nitrc.org/plugins/mwiki/index.php/neurobureau:INDIpreproc. The pixel-level mIF dataset is published with Li. et al [57]. The cell-level mIF dataset is publicly available through Mendeley Data https://data.mendeley.com/datasets/mpjzbtfgfr/1.

## 8 Code Availability

Code for data analysis and simulation in this manuscript is available at https://github.com/JiangmeiRubyXiong/PAMS

