## Supplemental Figures for "Facilitating Valid Statistical Inference in Biomedical Image Synthesis"

Supplementary Figures

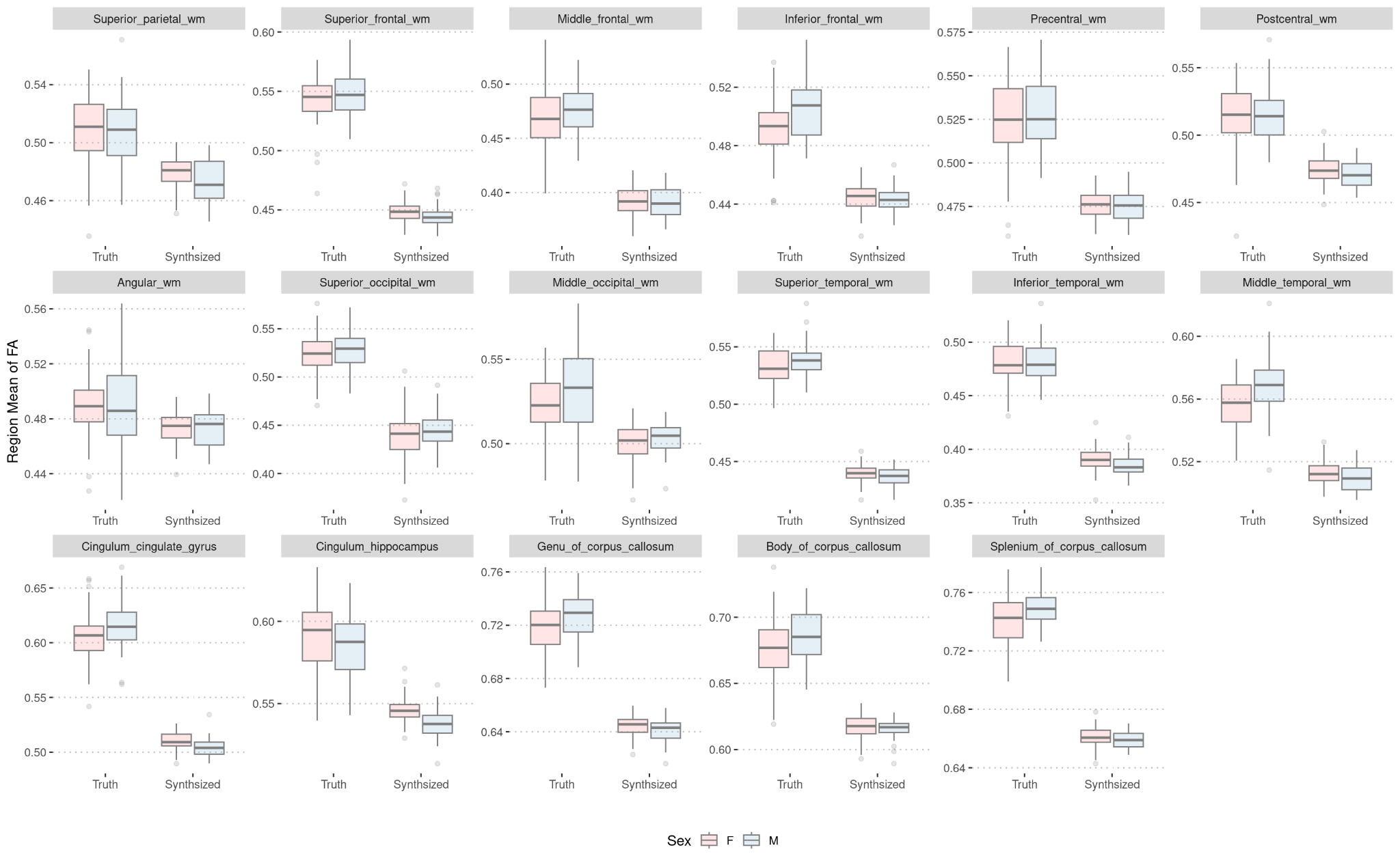

Figure S1. Comparison of all regions for neuroimaging. Each panel represents a white matter region. The values on the y-axis are the regional mean of FA and the two columns on the x-axis are the type of values (truth, synthesized), separated by sex.

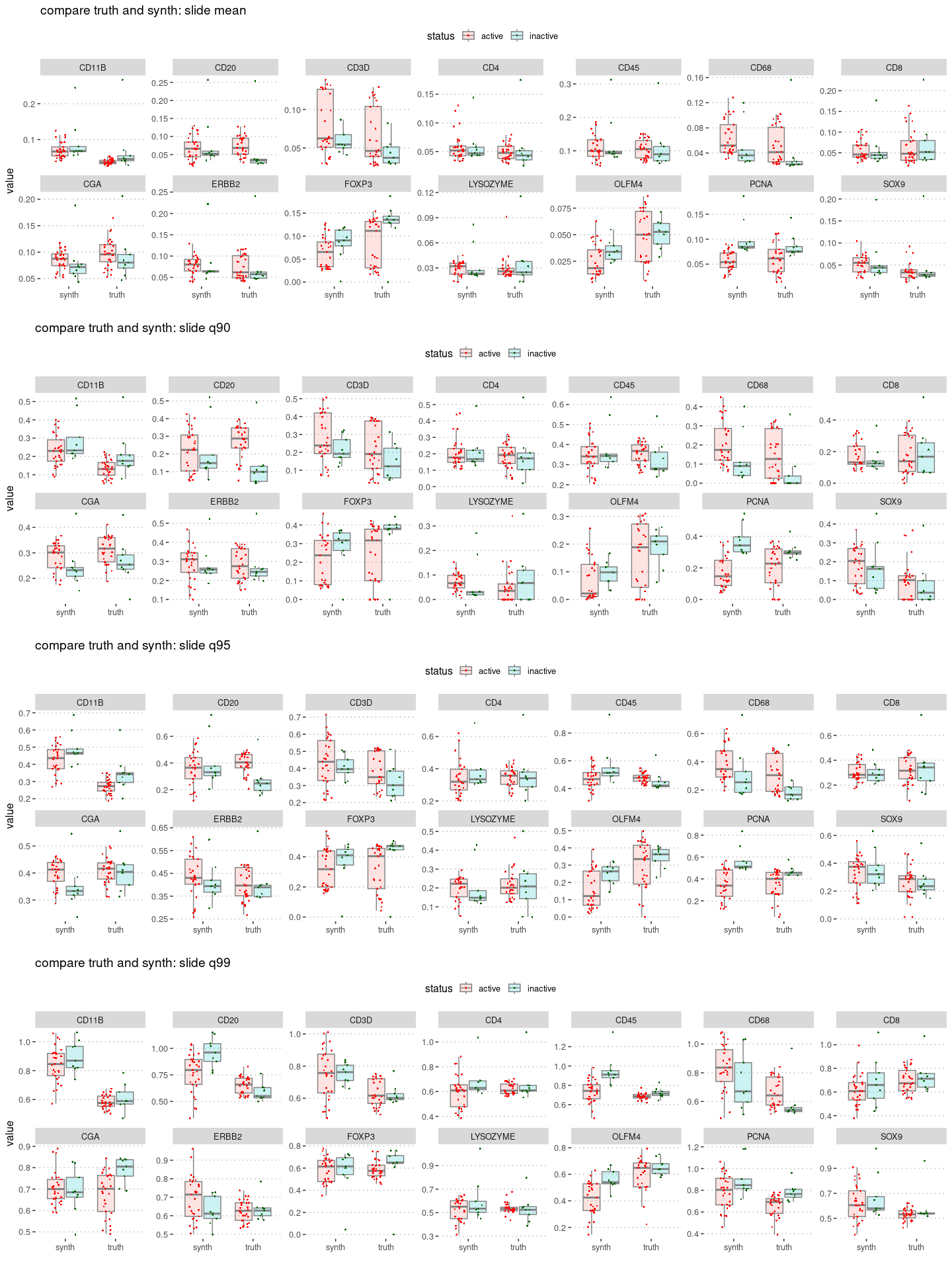

Figure S2. Comparison of all markers for mIF image level data, for mean, 90th, 95th, 99th quantile. Each panel represents a marker for a certain percentile. The values on the y-axis are the quantile value of pixel intensity expressions, and the two columns on the x-axis are the type of values (truth, synthesized), separated by disease status (active, inactive).

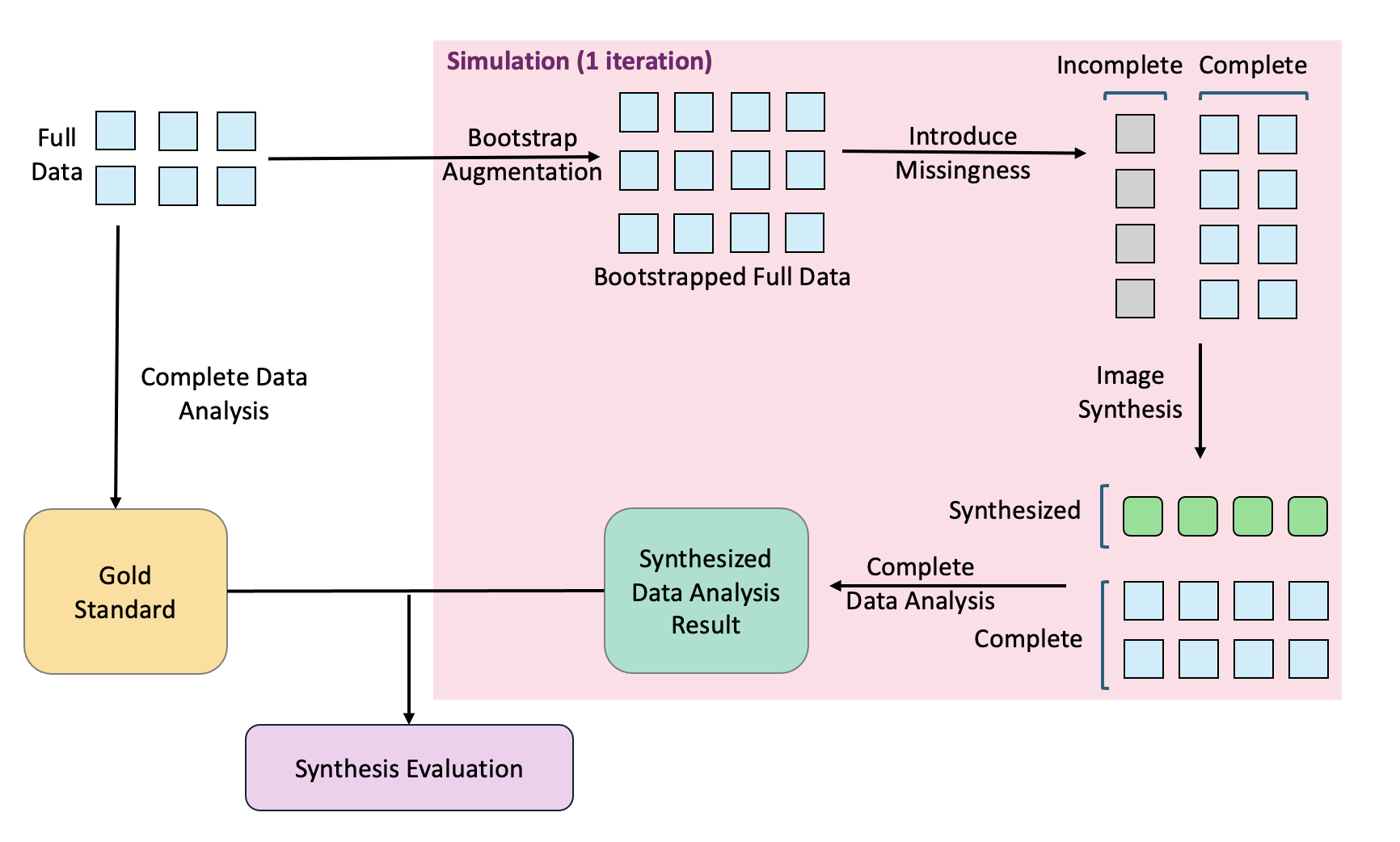

Figure S3. Diagram of simulation for the work flow of image synthesis. The gold standard is created by directly applying statistical analysis to the full data. Then for each iteration in the simulation, the full data is bootstrapped. Missingness is introduced to part of the data, and the designated missing images are synthesized using the model trained by external data. The same statistical analysis is applied to the synthesized and complete images, generating the synthesized data analysis result. Finally, the synthesized data analysis results from all simulation, along with the gold standard, goes into the synthesis evaluation.

# a)

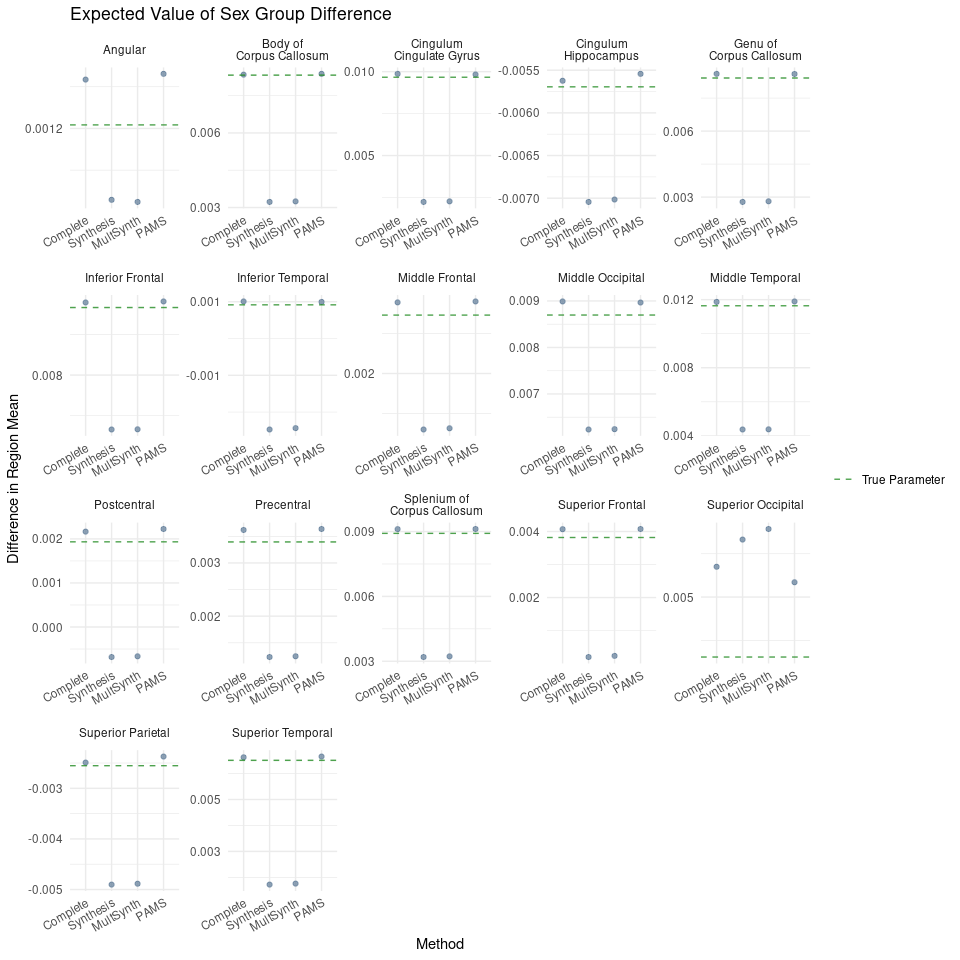

b)

c)
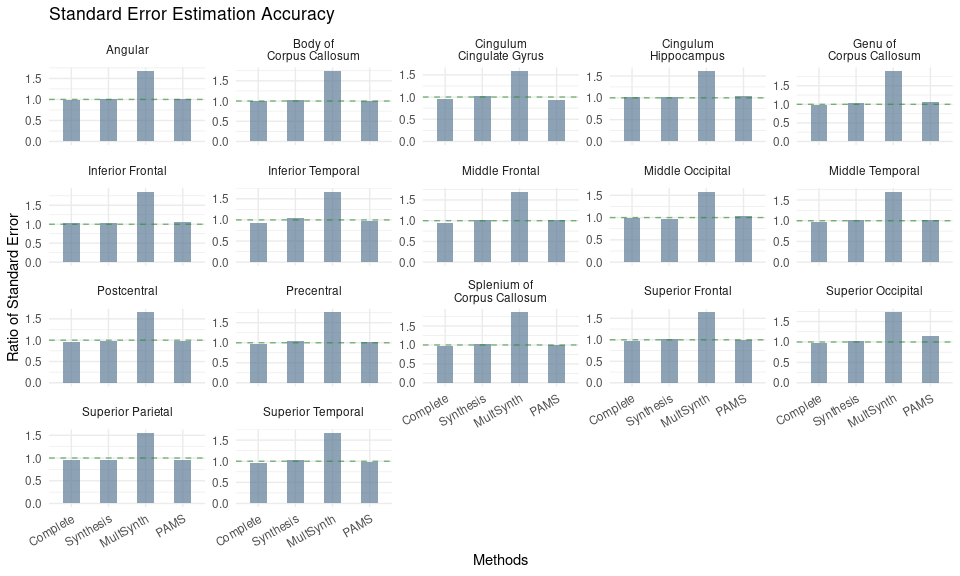

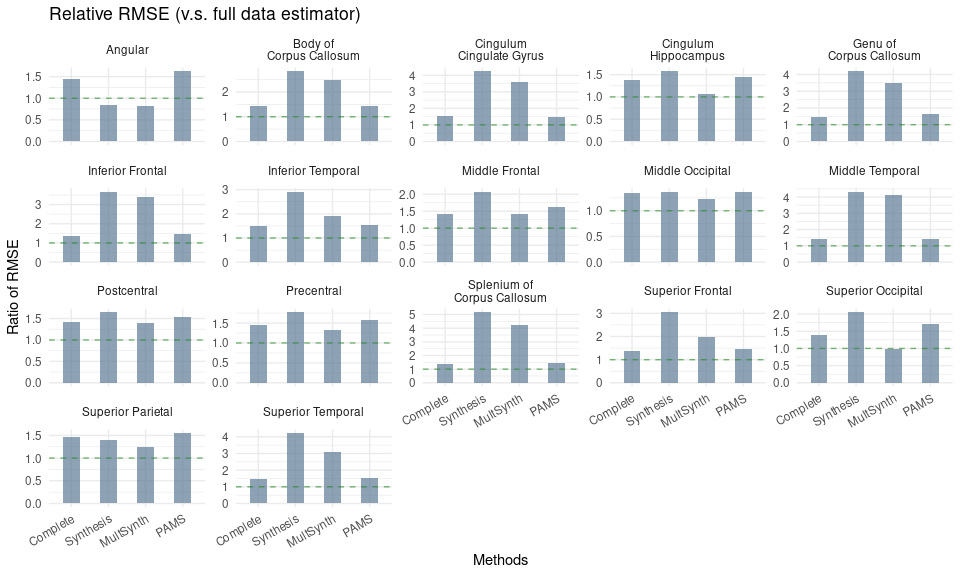

(d)
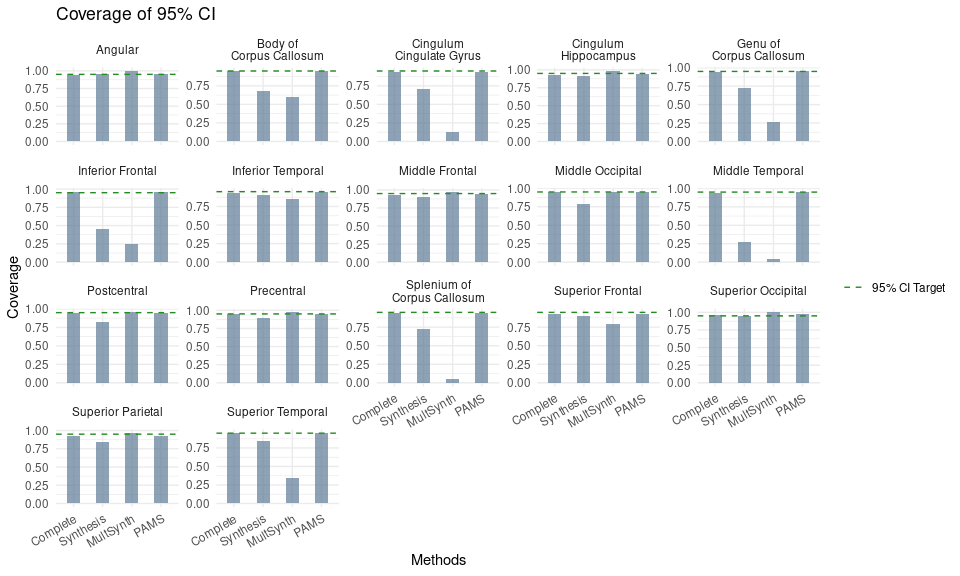

Figure S4. Bootstrap analysis of sex differences in fractional anisotropy (FA) in white matter regions comparing complete case analysis (“Complete”) with synthesis analysis methods. Results are shown for comparing GAN-based image synthesis (“Synthesis”), and Phenotype-Aware Multiple Synthesis using the same model (“PAMS”).

a)
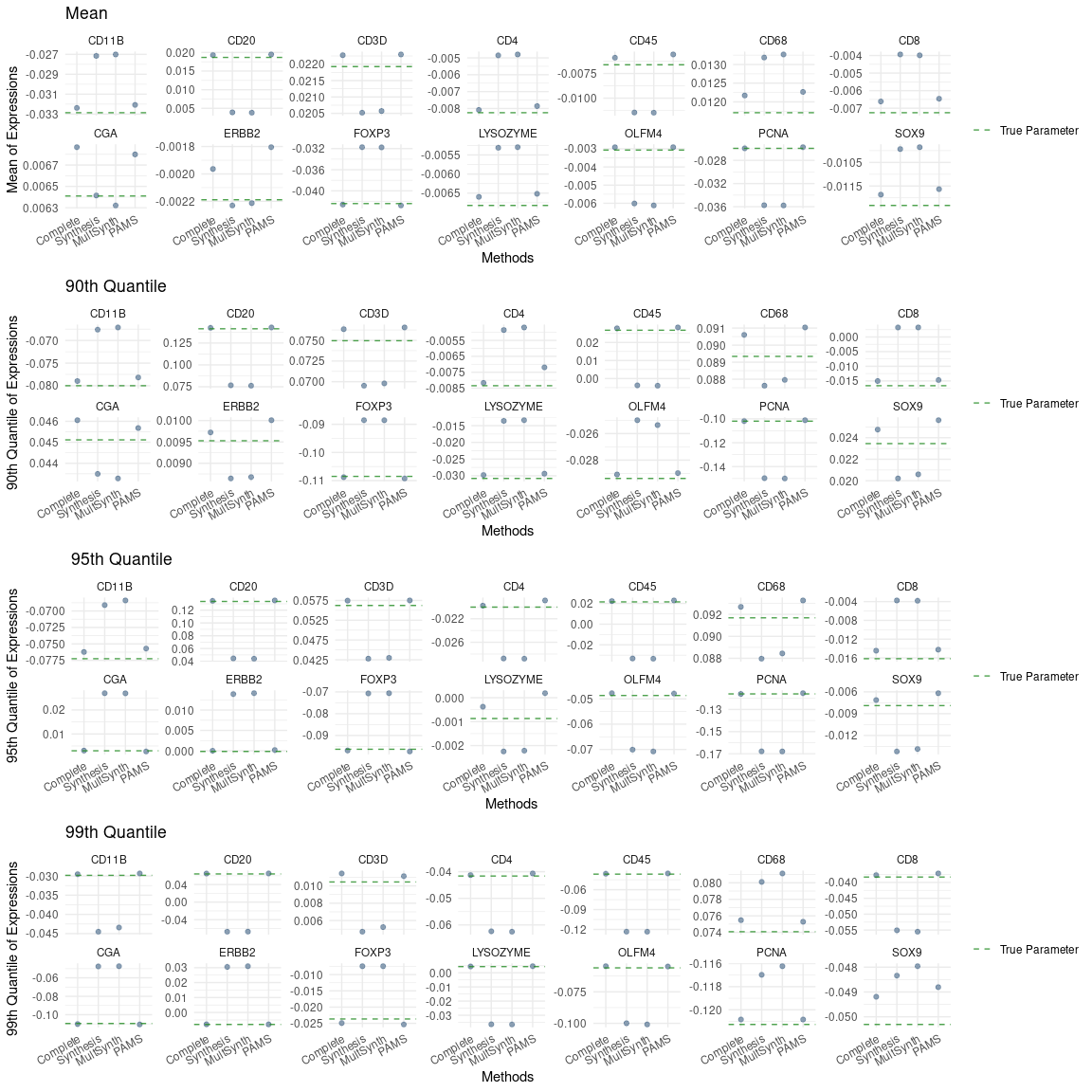

b)
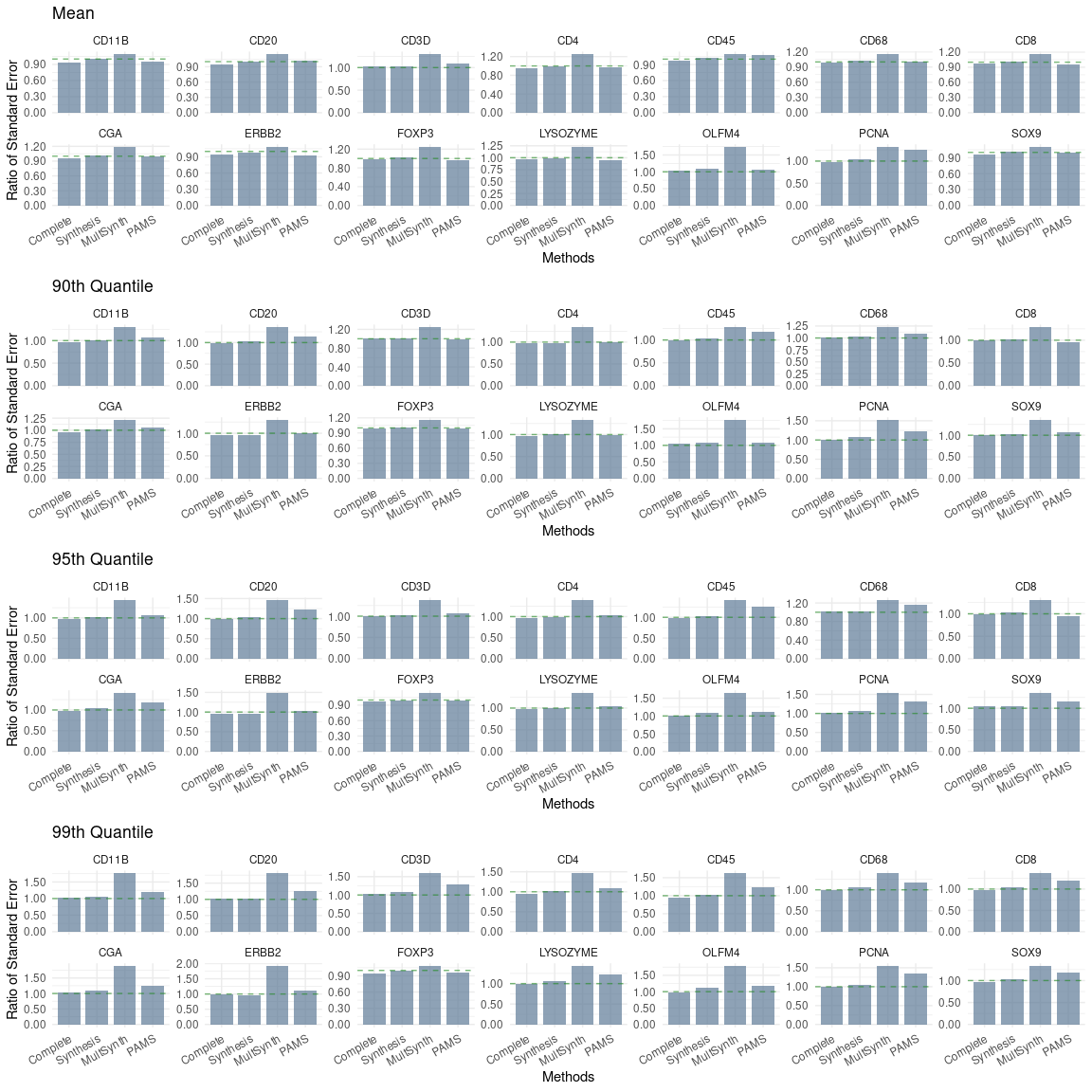

c)
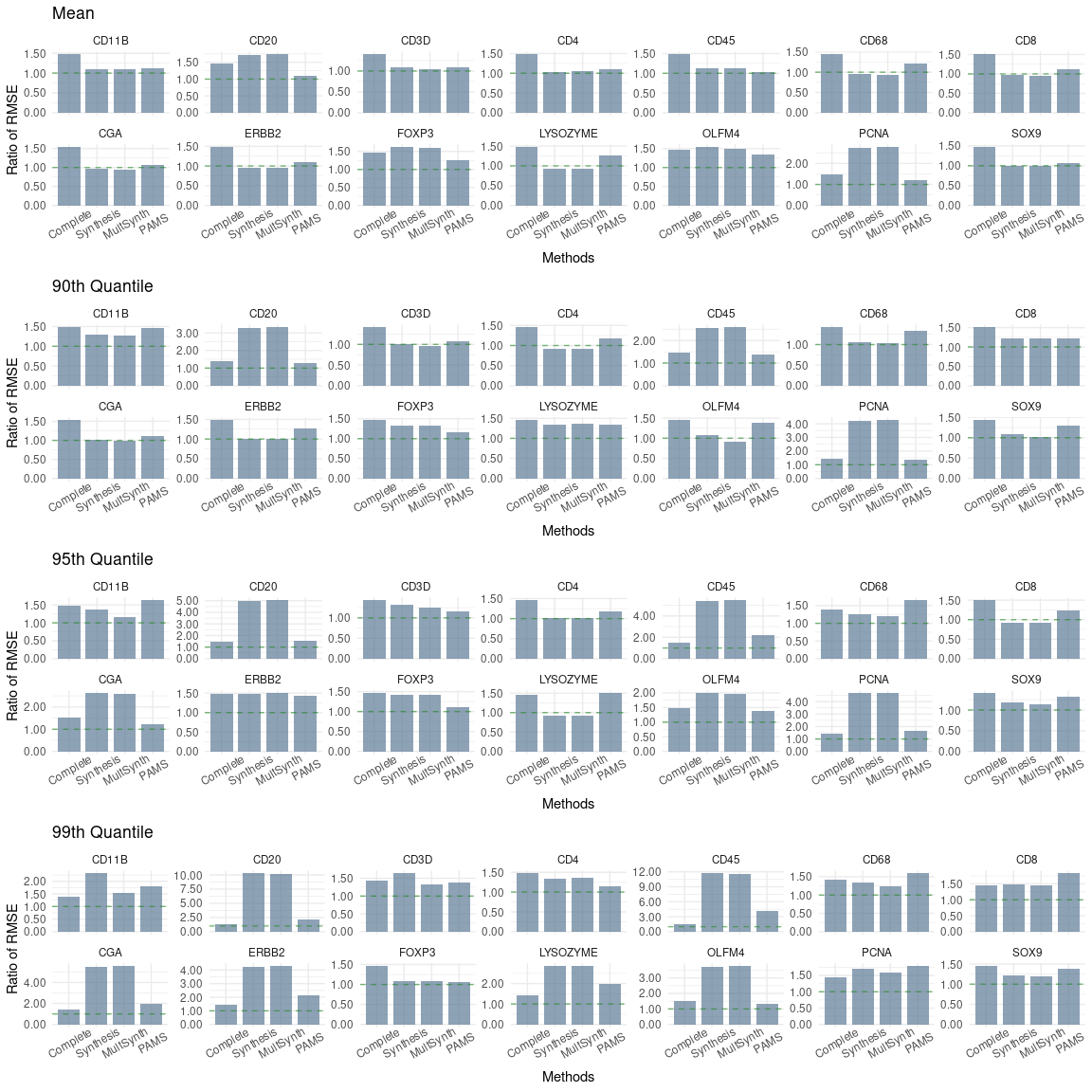
d)
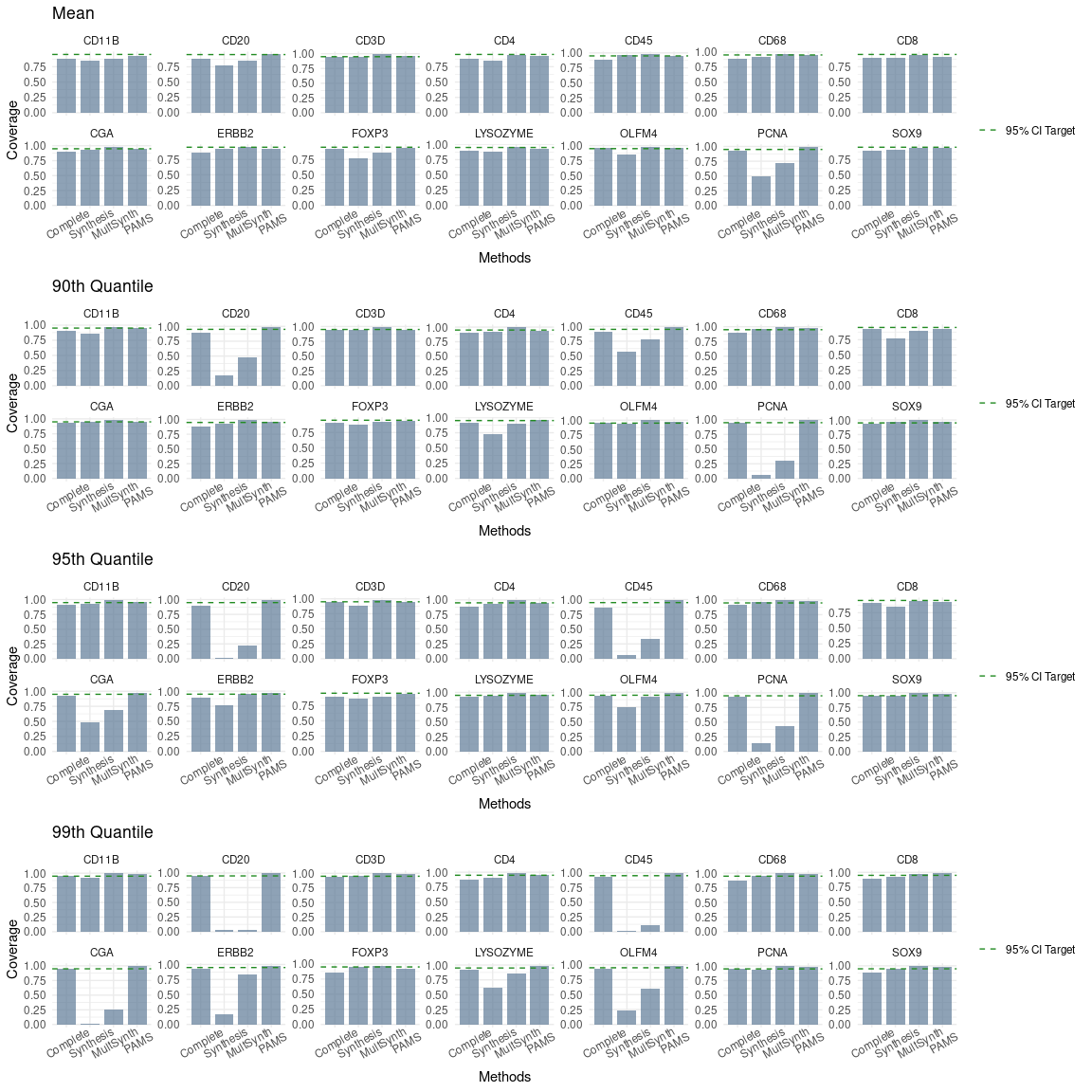

Figure S5. mIF image-level bootstrap analysis of disease status differences (active versus inactive) in the mean, 90th, 95th and 99th percentile expression comparing complete case analysis (“Complete”) to the synthesis analysis methods (“Synthesis” and “PAMS”) for all key markers.

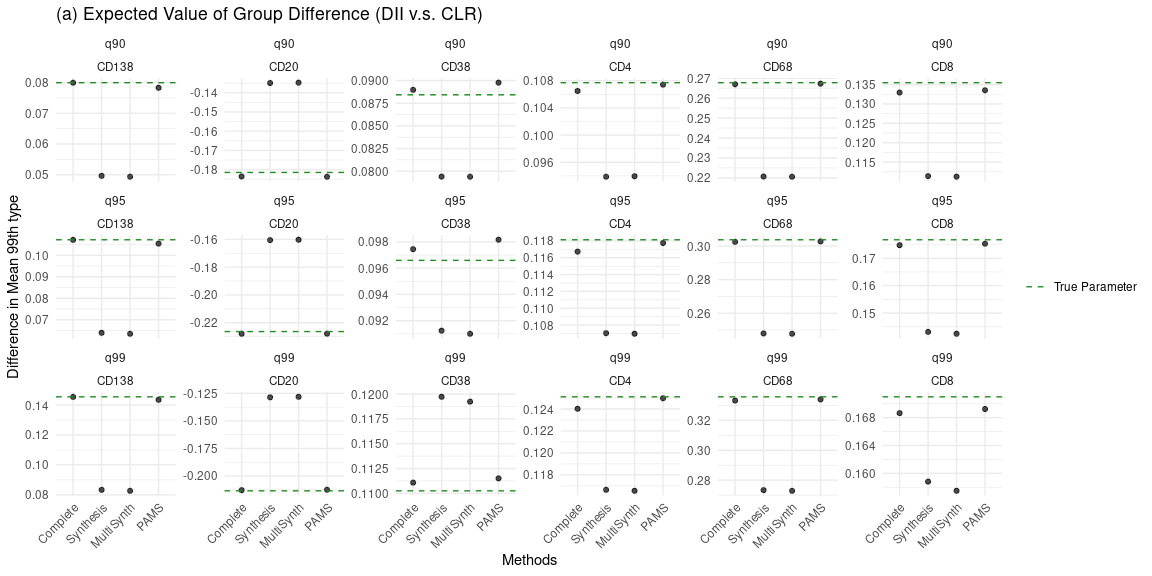

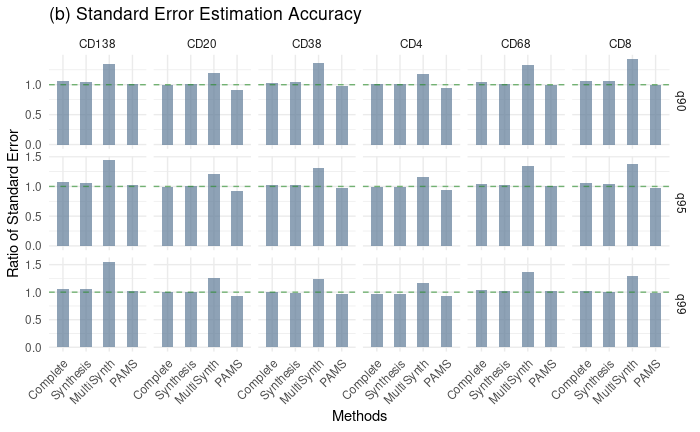

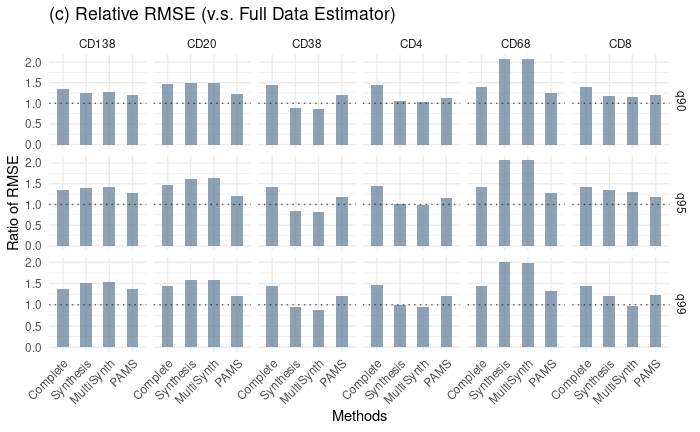

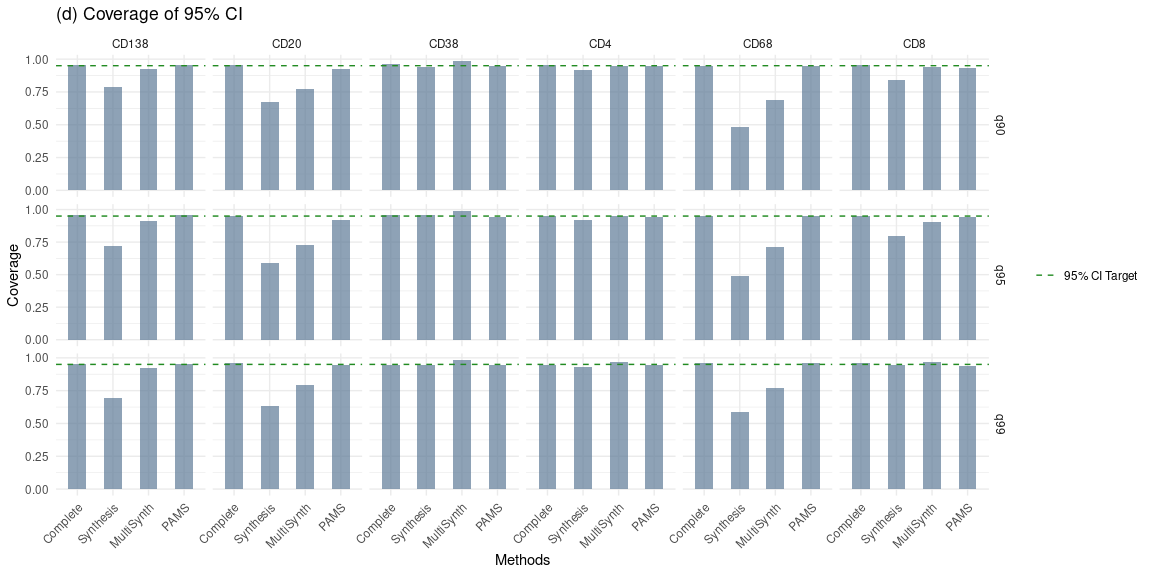

Figure S6. mIF cell-level bootstrap analysis of cancer type differences (CLR vs. DII) in the 99th percentile expression comparing complete case analysis (“Complete”) to the synthesis analysis methods (“Synthesis” and “PAMS”) for all key markers.
